# Interaction dynamics between epithelial cysts captured by tissue rheology

**DOI:** 10.1101/2025.08.11.669628

**Authors:** Marie André, Linjie Lu, Michèle Lieb, David Gonzalez-Rodriguez, Daniel Riveline

## Abstract

Epithelial cysts are minimal structures involved in morphogenesis. They are fluid-filled cavities surrounded by an epithelial monolayer. Cysts grow during development and their interactions shape organs. While their growth dynamics as single structures are well characterised, the physical mechanisms underlying their interaction remain poorly understood. Here we design a minimal assay of interacting cyst doublets based on microfabrication, quantitative biology, and theory to show that Madin-Darby Canine Kidney (MDCK) cyst interactions are essentially determined by the rheological properties of their epithelial monolayers. We report two phases of interaction between epithelial cysts: coalescence of cellular monolayers and lumen fusion, with similar speeds of 0.3 µm/h. We modulate the distribution of interaction phenotypes by reducing cell-cell adhesion using E-cadherin knock-out MDCK cells and we report that E-cadherin depletion promotes lumen fusion. Remarkably, the dynamics of coalescence and fusion are conserved between both cell lines. To understand the conserved speeds and the effect of cell-cell adhesion, we model the mechanical behavior of cyst doublets as a complex fluid to predict a speed determined by viscosity, a stretch-dependent monolayer tension, and the adhesion energy between cells. We measure these parameters through rheological experiments using micropipette aspiration and lumen drainage, which span the full range of stretch. A key insight from this analysis is that accounting for the tension’s dependence on stretch is is essential to capture the different dynamics observed during cyst interaction. Using these rheological measurements, we successfully recapitulate the conserved speed. Altogether, our results open new perspectives to understand tissue dynamics during organogenesis through simple physical arguments.

**Significance Statement:** Cysts are fluid-filled cavities sur-rounded by a cellular monolayer. Interactions between cysts are important during morphogenesis, yet their governing rules are poorly understood. Here we designed a doublet assay using microfabrication to control cyst geometry and characterized their interaction dynamics. We report that speeds of coalescence and fusion between epithelial cysts are conserved even for cell lines depleted in E-cadherin, a central adhesion protein. We measured their rheological parameters by two independent methods, micropipette aspiration and laser cutting. Using a simple model and scaling arguments, we demonstrate that these parameters are sufficient to explain interaction speeds. This study sheds light on the interplay between tissue dynamics and rheology and its implication in orchestrating morphogenetic events *in vitro* and *in vivo*.

---

**F**rom the first stages of embryogenesis to the formation of functional organs, cells proliferate, polarize, generate fluid-filled cavities surrounded by cell layers, the lumens (1–7). The cyst can be viewed as a basic structure – a lumen enclosed by an epithelial monolayer (1). Whereas the dynamics of organogenesis can result from diverse mechanisms, the formation of cysts is a conserved step.

Beyond their emergence, cysts interact, fuse and are remodelled over time. These dynamics eventually lead to the generation of tubes along which liquid can be secreted by cells, allowing circulation of materials within organs and across the body (8–10). In other cases, large cyst formations arise, exhibiting a variety of shapes in the colon or intestinal tract (6), for example.

Recently, rheological measurements have been reported as essential to explain morphogenesis. *In vivo*, viscosity measurements enabled the physical characterisation of Zebrafish early embryogenesis (11, 12). Similarly, during somitogenesis, viscosity gradients across the body axis were reported to correlate with elongation (13). Rheology is thus central to capture tissue dynamics. In this context, interaction between epithelial cysts also requires characterisation integrated with theory to eventually uncover the physical principles underlying their dynamics. How these spherical structures interact and how their behaviour is influenced by tissue-scale mechanical properties remain to be elucidated.

Here, we combine microfabrication, rheological modelling, and quantitative measurements to investigate this question. Using the Madin-Darby Canine Kidney (MDCK) epithelial cell line, we design a new setup – the *cyst doublet assay* - to track and characterize the spatio-temporal interactions between cysts and the eventual fusion of their lumens over up to 12 days. We probe the effect of cell-cell adhesion on the distribution of interaction phenotypes and we find that E-cadherin depletion promotes lumen fusion, while the interaction dynamics and speed remain similar between wildtype (WT)-WT and E-cadherin knock-out (E-cad KO)-E-cad KO cyst doublets. To rationalize our experimental observations, we model the physics of cyst interaction within a viscoelastic fluid framework. A key idea in this model is that, unlike surface tension in liquids, cyst monolayer tension combines an adhesion component with an elastic component that depends on the degree of stretching, as previously proposed (14, 15). This framework links the interaction dynamics to the rheological properties of cysts. To parametrize and test the model under different adhesion levels, we measure the mechanical properties of both MDCK WT and E-cad KO cysts through micropipette aspiration and laser cutting experiments, which enables us to determine viscosity, elastic tension and adhesion energy of cysts. Using these measurements, we successfully recapitulate the coalescence and fusion dynamics.

## Results

### Interaction of MDCK cysts occurs through monolayer coalescence and lumen fusion: the doublet assay

In order to study and track interactions between cysts of similar dimensions in a reproducible manner, we designed a new microfabricated assay (Fig. 1). We trapped single MDCK cells in cavities (see Materials and Methods) (16, 17). We incubated them with a 3D matrix (Matrigel), which triggered the growth of cysts over time (Fig. 1B). The distance between cavities was designed such that two neighbouring cysts eventually met at about 3-4 days after plating, considering the proliferation rate and the increase in cyst size over time (Supplementary Fig. 1).

**Fig. 1.**
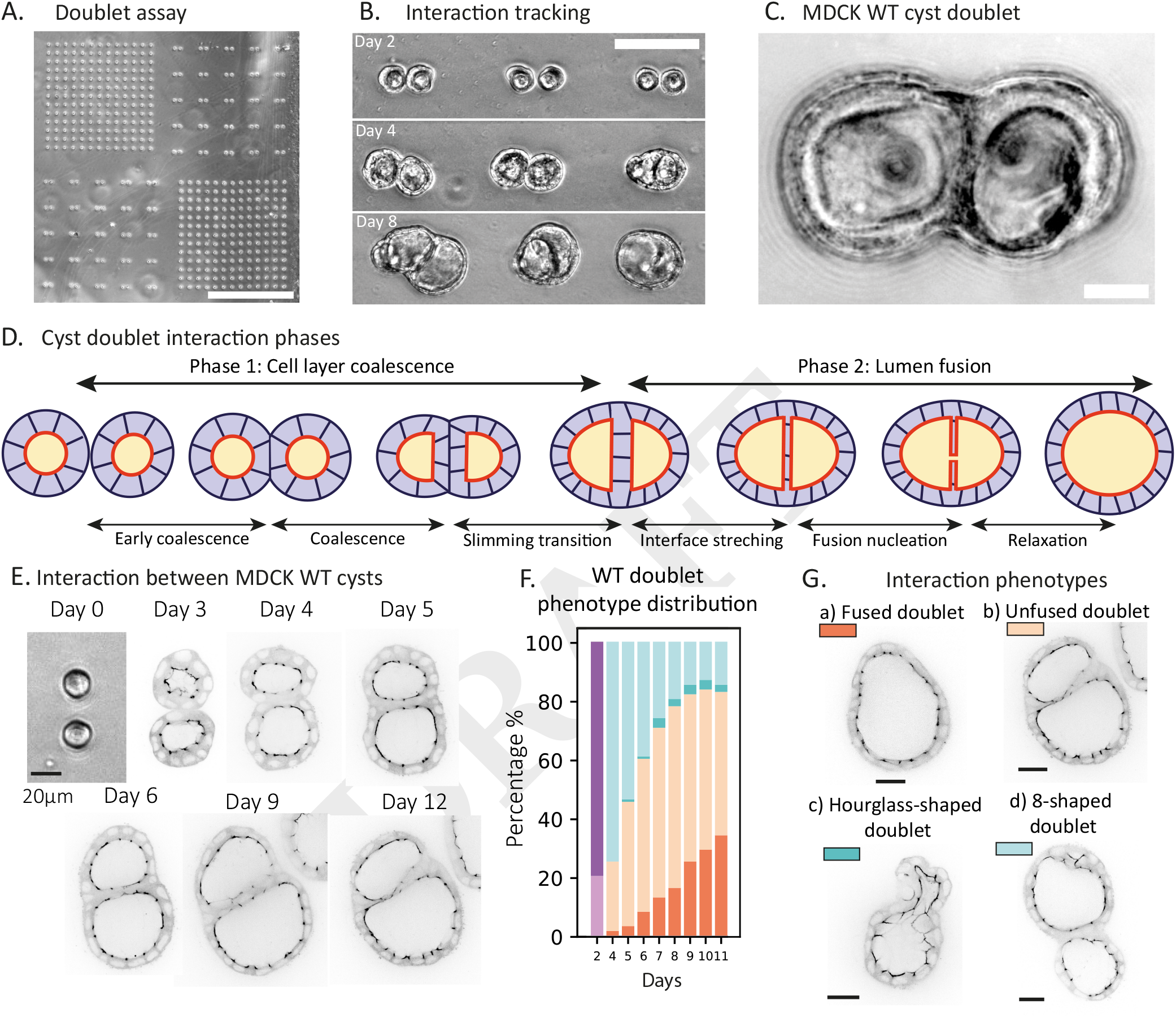
Interaction of MDCK cysts occurs through coalescence and fusion of lumens with four phenotypes. A) Cavities used to track doublets during 8 days and beyond. Cavities have a diameter of 17 µm and the inter-distance between two cavities is 15 µm. Scale bar 500 µm. B) The sample chessboard design enables us to identify each doublet and perform the tracking efficiently. Scale bar 100 µm. C) Doublets of MDCK cysts interacting at day 7 after seeding. Cavities have a diameter of 17 µm to ensure each cyst starts from a single cell. Scale bar 20 µm. D) Schematics of the interaction between cysts. This is a two-phase process featuring the cell layers deformation and coalescence (Phase 1) followed by cellular reorganization and lumen fusion (Phase 2). Phase 1 is composed of an early coalescence lasting for 5-10 hours after the beginning of interaction, a longer coalescence. Before the initiation of a hole in the interface leading to lumen fusion, we observe several cellular dynamics including cell interaction in the neck region and interface thinning. E) Interaction between two MDCK cysts expressing fluorescent ZO1-mNeonGreen. Coalescence occurs between day 3 and 5. Scale bar 20 µm. F) Quantification of phenotypes observed during the interaction of MDCK WT cysts. Purple colours at day 2 correspond to the percentage of cyst doublets in contact (light purple) and not in contact (dark purple). The other colours correspond to each phenotype presented in G. G) The four phenotypes present on interacting samples. The outer shape of doublets can be either completely coalesced (top line in orange, a. and b.) or incompletely coalesced (bottom line in blue, c. and d.). The inner luminal structures can be either fused (dark colours, left column, a. and c.) or unfused (light colours, right column, b. and d.). Colours correspond to the phenotype distribution in F. Scale bar 20 µm.

Our assay enabled us to follow cyst doublets over 12 days (Fig. 1E) and their spatial organizations in a “chess-board” layout allowed us to increase the acquired statistics within each experiment (Fig. 1A). Tracking cysts enabled us to distinguish two phases of interaction, coalescence of monolayers and fusion of lumens (see schematics Fig. 1D). The interaction proceeded together with local cellular rearrangements, including slimming transitions from two monolayers between cysts to a single one and thinning of the interface between lumens (Fig. S2F,K).

### Interactions of MDCK cysts display four phenotypes

The observation of various phenotypes at the same day of culture prompted us to define carefully the signature of interacting doublets (Fig. 1C,G). We identified four types of doublet shapes: coalesced cysts with quasi-spherical outline and fused lumens (fused doublet, Fig. 1Ga, dark orange); coalesced cysts with quasi-spherical outline but separated lumens (unfused doublet, Fig. 1Gb, light orange); incomplete coalescence and fused lumens (hourglass-shaped doublet, Fig. 1Gc, dark blue); incomplete coalescence and non-fused lumens (8-shaped doublet, Fig. 1Gd, light blue). We parametrized these shapes and reported that their respective signatures were indeed distinct (Fig. S2A,E). Finally, we quantified their distributions and showed that the percentage of each phenotype evolves over time. Half of the cyst doublets are completely coalesced at day 5 and 83% are completely coalesced by day 11. When considering the final state of the interaction, we find that 49% of doublets are unfused, 34% are fused, 15% are 8-shaped, and 2% are hourglass-shaped (Fig. 1F). These distributions suggest that cell monolayer coalescence and lumen fusion can be seen as parallel events whose outcomes are not directly determined by each other.

### Reduction in cell-cell adhesion facilitates fusion

In order to gain insight into the mechanisms underlying the interaction of cysts, we systematically compared MDCK WT cysts with MDCK E-cad KO cysts depleted for adherens junctions (18). We hypothesized that modified levels of adhesion between cells would impact the dynamics of cyst interaction.

When considering phenotype distribution, E-cad KO cyst interaction showed a similar percentage of completely coalesced phenotypes compared to WT cyst interaction (unfused and fused doublets, compare Fig. 2A-B with Fig. 1E-F). However, by day 11, 89% of E-cad KO doublets had fused their lumens, compared to 34% of WT doublets (Fig. 2C).

**Fig. 2.**
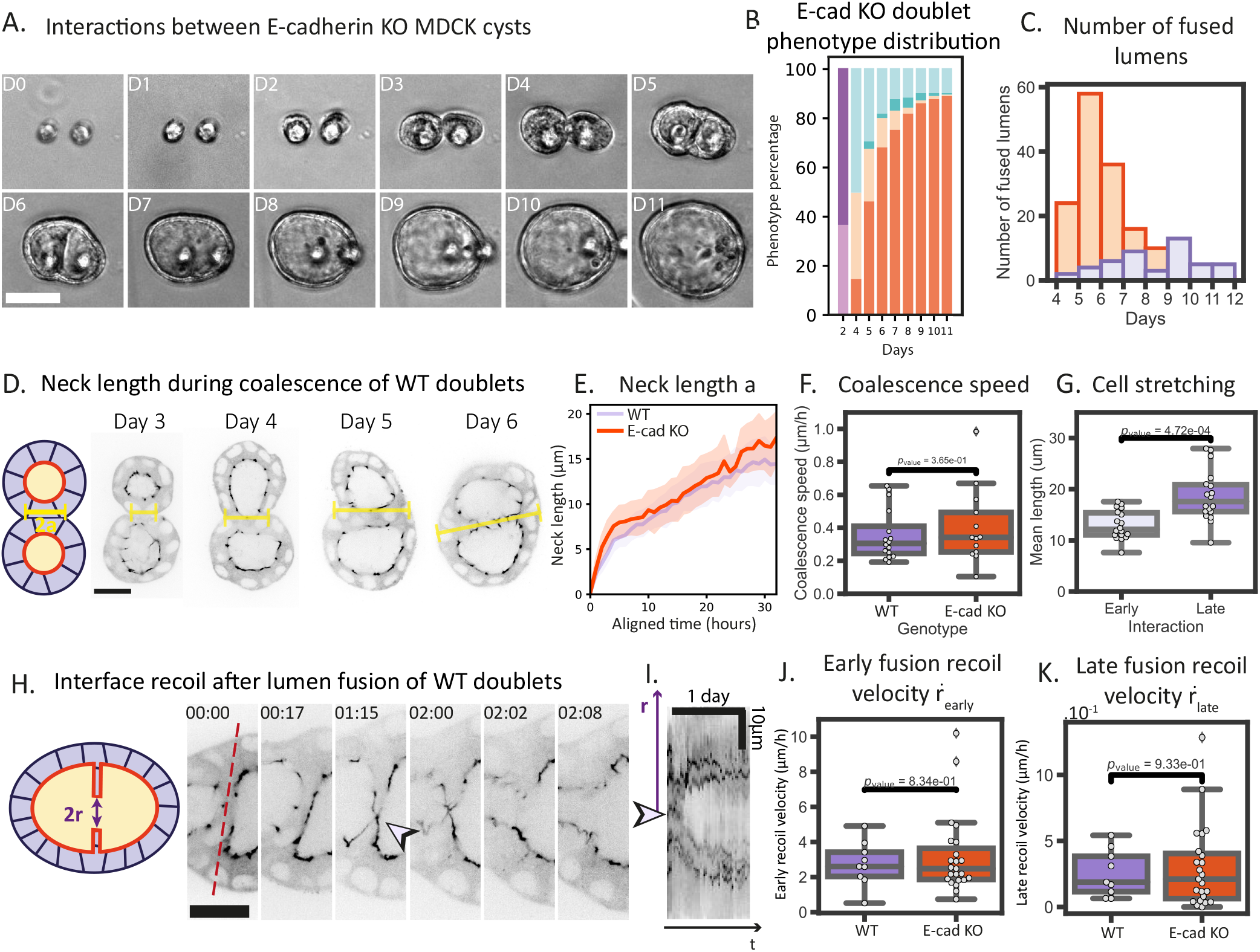
Interaction dynamics is captured through coalescence velocity, fusion velocity and fusion distribution for interaction of MDCK WT and E-cad KO cysts. A) Tracking over 12 days of interaction between two MDCK E-cad KO cysts starting from single cells. Scale bar 50 µm. B) Percentage of phenotypes observed in interactions between E-cad KO cysts (N=3, n=168). Colour code corresponds to phenotypes presented in Figure 1F-G. C) Number of new fused lumens over days of interaction. Interactions between MDCK WT cysts are indicated in purple and MDCK E-cad KO ones in orange. (N_WT_=3, n_WT_=44, N_KO_=3, n_KO_=150). D) Neck length measurement on interacting coalesced MDCK WT cysts doublets expressing ZO1-mNeonGreen. Scale bar 20 µm. E) Neck length a measured on 40 hours time-lapse data aligned on the beginning of interaction. Purple curves indicate WT doublets (N_WT_=4, n_WT_=16) and orange ones E-cad KO doublets (N_KO_=3, n_KO_=14). WT and E-cad KO doublets display the same interaction dynamics. Curves indicate means ± 95% confidence interval. F) Comparison between MDCK WT and MDCK E-cad KO cysts doublets coalescence speed (N_WT_=4, n_WT_=16, N_KO_=3, n_KO_=14). G) Average cell stretching during early and late interaction of MDCK WT cysts (N=3, n=18). H) Fusion of lumens MDCK WT cysts doublets expressing ZO1-mNeonGreen aged 4 days after seeding. The red dashed line indicates the line along which the kymograph was generated in I. The arrow indicates the position of the initiation of the lumen’s fusion. Time in days:hours, scale bar 20 µm. Movie S2. I) Kymograph of the lumen fusion presented in H. The darker lines correspond to the interface recoil. The large arrow indicates the position for initiation of lumen fusion. The smaller vertical and horizontal ones indicate respectively the axis along which the recoil radius *r* and the time *t* are read. J) Fusion early recoil velocity 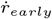 in WT and E-cad KO doublets measured on timelapse data (N_WT_=5, n_WT_=9, N_KO_=4, n_KO_=23). K) Fusion late recoil velocity 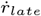 in WT and E-cad KO doublets measured on timelapse data (N_WT_=5, n_WT_=9, N_KO_=4, n_KO_=23).

Interaction phenotype distribution is impacted by the reduction of cell-cell adhesion. We thus refined the quantification of interaction dynamics and tested whether monolayer coalescence and lumen fusion dynamics were dependent on cell-cell adhesion.

### Coalescence and fusion speeds do not depend on cell-cell adhesion

To capture the coalescence dynamics, we quantified the neck length elongation at the contact between two cysts over several hours of their interaction (Fig. 2D-G, Movie S1). We compared the neck length increase between MDCK WT cyst doublets and MDCK E-cad KO cyst doublets and we obtained similar velocities: 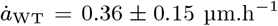 for WT cyst doublets and 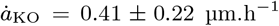 for E-cad KO cyst doublets (Fig. 2F). This velocity constitutes a signature of the physical interaction of doublets. The similarity of coalescence speeds suggests that their dynamics are governed by a shared physical framework. We verified that the matrix did not significantly affect velocity by repeating the doublet assay with lower percentage of Matrigel (Fig. S3B-C). As the number of fused lumens was larger for E-cad KO cyst interactions, we further quantified lumen fusion dynamics by tracking retraction of the cellular layer between lumens during their fusion (Fig. 2H-K, Fig. S4, Movie S2). The measurements revealed that the early and longterm interface retraction velocities were conserved between WT and E-cad KO doublets. These early recoil velocities were 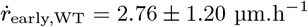 for WT and 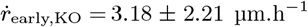 for E-cad KO (Fig. 2J) and the longterm ones were 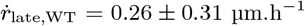 for WT and 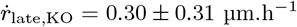 for E-cad KO (Fig. 2K).

The consistency between interaction dynamics suggests that coalescence and fusion are governed by common physical mechanisms. Drawing inspiration from previous work on cellular aggregates (19–24), we hypothesized that the dynamics of MDCK cyst coalescence and fusion are set by their rheological properties. We thus turned to rheological experimental characterization and modelling of cysts to investigate whether monolayer tension and viscosity could quantitatively account for the measured interaction dynamics.

### Mechanics of cysts are distinct between WT and E-cad KO cell lines as shown by micropipette aspiration and laser cutting

Due to its elastic properties, cyst monolayer tension *σ* increases as the cyst stretches (14, 15). The relationship can be expressed as *σ* = *W*_cc_ + *f* (*L/L*_base_), where *W*_cc_ is the cell-cell adhesion energy, and *f* (*L/L*_base_) is an elastic term that depends on cell stretching. Here, *L/L*_base_ represents by the ratio of the apico-basal cell size *L* to its baseline value *L*_base_ in a relaxed cyst. To quantify these two components, we combined two rheological techniques: micropipette aspiration (25), which measures viscosity *η* and monolayer tension *σ*_asp_ in stretched cysts (*L > L*_base_); and laser cutting, which provides the baseline tension *σ*_base_ at *L* = *L*_base_ and the surface tension of deflated cysts *σ*_cut_ ≈ *W*_*cc*_ for *L* ≪ *L*_base_.

We performed micropipette aspiration experiments along former assays (25–27) (Fig. 3A-B, Movies S3-4). Briefly, we determined the critical pressure needed to aspirate the monolayer which allowed us to estimate the monolayer tension under elastic stretching (Fig. 3A) of *L* ≈ 1.5*L*_base_ (Fig. S5). At larger aspiration pressures, we measured the velocity of monolayer entry into the pipette, which allowed us to determine the viscosity (Fig. 3B). We obtained pairs of values for MDCK WT and E-cad KO cysts: MDCK WT cysts had a monolayer tension *σ*_asp_ = 16 ± 4 mN.m^−1^ and a viscosity *η* = 2.4 ± 2.1.10^5^ Pa.s (purple on Fig. 3B,D), and MDCK E-cad KO cysts display a monolayer tension *σ*_asp_ = 7 ± 2 mN.m^−1^ and a viscosity *η* = 0.8 ± 0.5.10^5^ Pa.s (orange on Fig. 3B,D). Both monolayer tension and viscosity were decreased in MDCK E-cad KO cysts. Ratio between viscosity and tension gives a speed, which is strikingly constant between cell lines, i.e., about 0.1 µm.s^−1^ (Fig. S5A). This result echoes the similar speeds measured for coalescence and fusion. However, the speed obtained by scaling is three orders of magnitude faster. This discrepancy likely arises because the monolayer is under elastic stretching during micropipette aspiration, whereas during coalescence, the dynamics are driven by the adhesion term alone and stretching plays no role (see also cell elongation during coalescence outlined in Fig. S6).

**Fig. 3.**
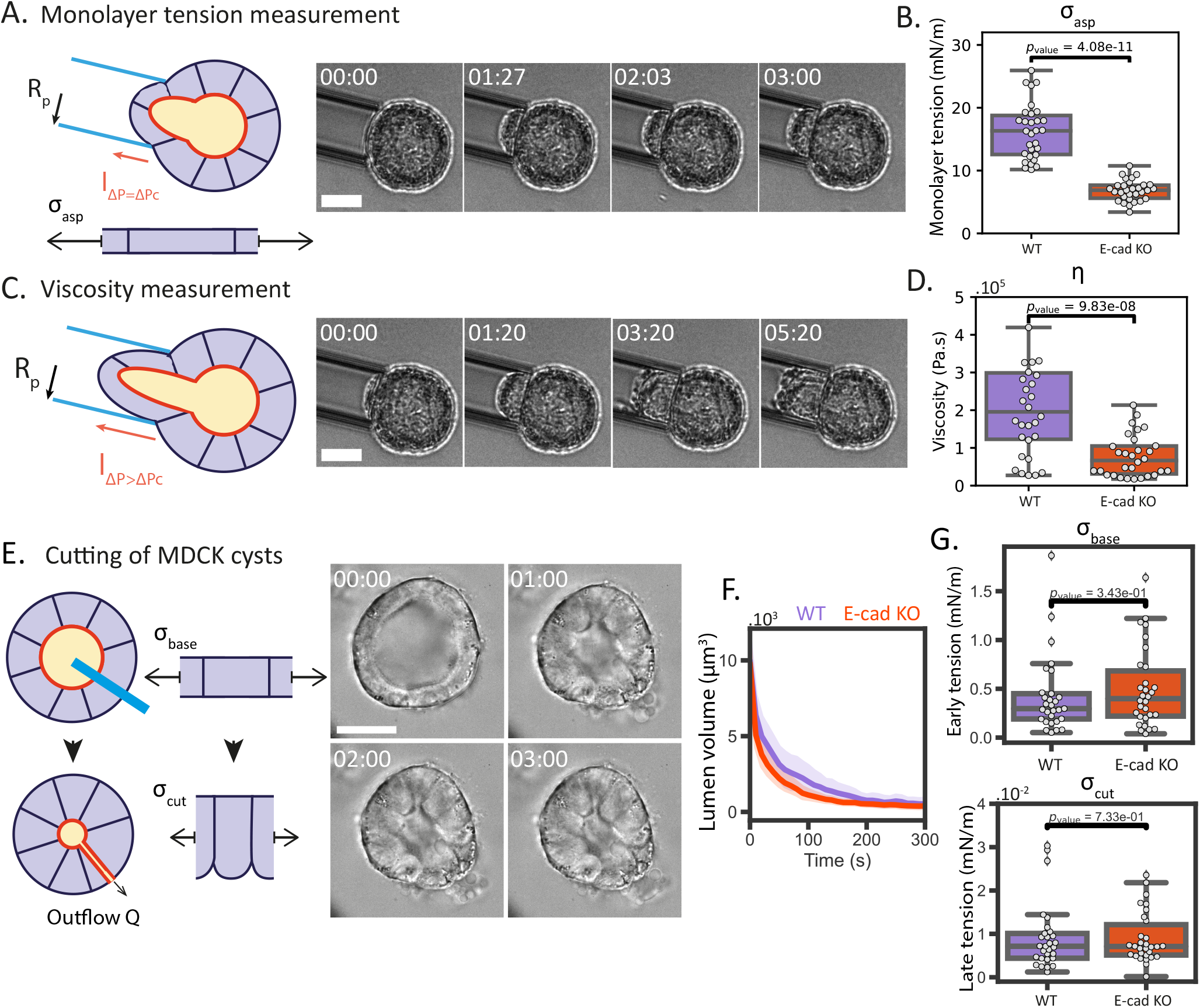
Mechanical properties of MDCK cysts through micropipette aspiration and laser cutting. A) Left: Schematics of micropipette aspiration to measure surface tension of cysts. The monolayer is stretched during aspiration as highlighted by the schematics giving rise to a tension *σ*_asp_. Right: Micropipette aspiration to measure surface tension of MDCK cysts aged day 4 after seeding. Time in mm:ss, scale bar 20 µm. Movie S3. B) Monolayer tension *σ*_asp_ measurements from micropipette aspirations for MDCK WT and E-cad KO cysts at day 4 after seeding (N=3, n=30). C) Left: Schematics of micropipette aspiration to measure viscosity of cysts. Right: Micropipette aspiration to measure viscosity of MDCK cysts aged day 4 after seeding. Time in mm:ss, scale bar 20 µm. Movie S4. D) Viscosity *η* measurements from micropipette aspirations for MDCK WT and E-cad KO cysts at day 4 after seeding (N=3, n=30). E) Left: Schematics of laser cutting to measure outflow Q, baseline tension and surface tension of cysts. The cut is performed on the blue line. The monolayer displays a baseline tension *σ*_0_ from which the elastic component is relaxed following laser cutting, giving rise to a relaxed tension *σ*_cut_. Right: Laser cutting of MDCK cysts aged day 4 after seeding to measure late surface tension. Time in mm:ss, scale bar 20 µm. Movie S5. F) Lumen volume time evolution after laser cutting for WT cysts (purple, N_WT_=3, n_WT_=28) and E-cad KO cysts (orange, N_KO_=3, n_KO_=38). G) Comparison of early (*σ*_base_, top) and late tension (*σ*_cut_, bottom) estimated from the cutting experiments between WT cysts and E-cad KO cysts (N_WT_=3, n_WT_=28, N_KO_=3, n_KO_=38).

To quantify this adhesion component, we performed lumen drainage through laser cutting experiments (28). In these experiments, short-term cyst emptying provided an estimate of monolayer tension at baseline cyst conditions (*L* ≈ *L*_base_), whereas long-term cyst emptying exhibited liquid-like relaxation as the elastic contribution of monolayer stretching dissipated (*L* ≪ *L*_base_, Fig. 3E,G and Fig. S5B-C for stretching). We thus performed laser cutting characterization of MDCK cysts. We report an early monolayer tension *σ*_base_ of the order of 0.42 ± 0.39 mN.m^−1^ for WT cysts and 0.50 ± 0.40 mN.m^−1^ for E-cad KO cysts. We measure a late monolayer tension *σ*_cut_ of the order of 0.0091 ± 0.0075 mN.m^−1^ for WT cysts and 0.0091 ± 0.0057 mN.m^−1^ for E-cad KO cysts (Fig. 3E,G, Movie S5). Using these latter values, which reflect purely adhesion-driven surface tension, a simple scaling argument yields a velocity estimate of the order of *σ*_cut_*/η* ≈ 0.1 µm.h^−1^, corresponding to the measured doublet coalescence and fusion speeds. With viscosity and tension terms now quantified, we proceeded to quantitatively rationalize the dynamics of doublet interaction beyond scaling arguments.

### Coalescence and fusion dynamics are recapitulated from rheological measurements

The dynamics of cyst interaction are captured by a minimal fluid model accounting for viscosity *η*, adhesion energy *W*_cc_ = *σ*_cut_, and a stretchdependent monolayer tension arising from elasticity. These parameters, quantified in rheological experiments, reflect the stage-dependent stretching of cells during interaction (Fig. 4A). Importantly, the three measured tensions correspond to distinct stages: *σ*_asp_ (aspiration) reflects the higher stretching during fusion nucleation; *σ*_base_ (early-time laser cutting) captures the baseline tension relevant for fusion recoil; and *σ*_cut_ (late-time laser cutting) measures the adhesion energy, which drives coalescence, since elastic stretching has largely relaxed by the end of the laser cutting experiment. This correspondence arises from the pairwise similarity in stretching ratios (*L/L*_base_) between rheological measurements and stages of interaction (Fig. 4A). This framework provides a quantitative description of coalescence, fusion nucleation, and lumen fusion.

**Fig. 4.**
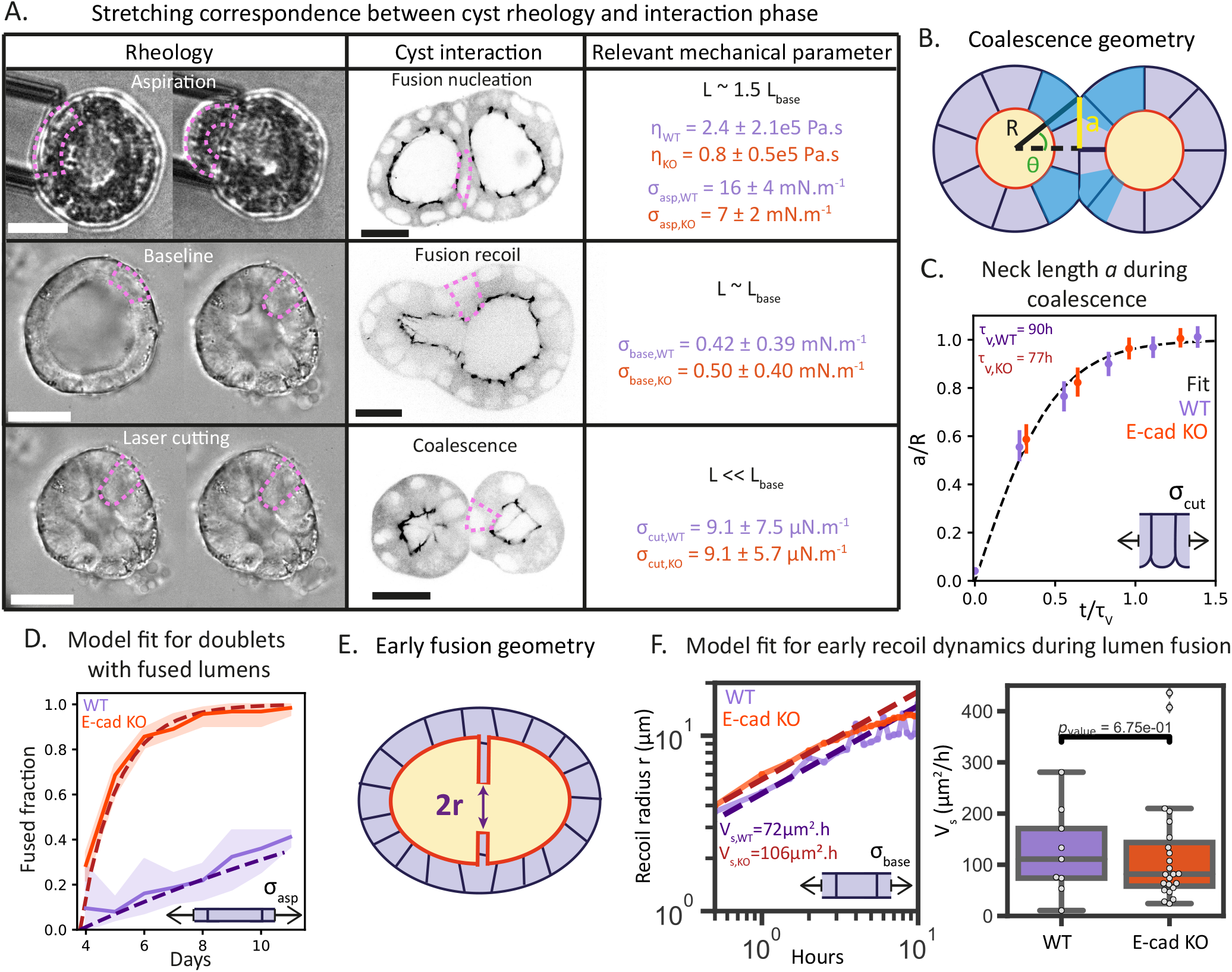
Comparison between theory and experiments. A) Table showing the correspondence of stretching levels (rows) between rheological experiments and interaction phases together with measured parameters (columns). On experimental images, the stretching of the monolayer is outlined in dashed pink. Scale bars 20 µm. B) Schematics of the geometrical description of cyst coalescence. *R* is the cyst radius, *a* the neck radius and *θ* the coalescence angle. The blue regions indicate cells that concentrate the viscous dissipation. C) Coalescence dynamics on the neck length data for MDCK WT and E-cad KO cysts interaction (respectively purple and orange). Neck radius *a* is normalized by mean cyst radius *R*, highlighting the similar interaction dynamics. The schematics insert illustrates the relaxed stretching state of the monolayer and the associated tension *σ*_cut_. The black dashed line corresponds to the adhesion-driven coalescence approximated solution for the estimated viscoelastic timescale *τ*_v_ fitted on the experimental data. The fitted characteristic times give *τ*_v,WT_ = 90 h and *τ*_v,KO_ = 77 h (SI for details on the fit). D) Predicted evolution of the fraction of fused doublets compared to the total coalesced doublets. The experimental data for MDCK WT and E-cad KO doublets (respectively purple and orange circles) are compared to prediction of the model for number of lumen fusion over days for Γ_WT_ = 0.06 day^−1^ and Γ_KO_ = 0.8 day^−1^ (SI for details on the fit). The schematics insert illustrates the stretched state of the monolayer and the associated tension *σ*_asp_. E) Schematics of the geometrical description for early fusion geometry. The recoil radius *r* increases over the fusion process. F) Left: Model fit for early recoil radius *r* of the interface during lumen fusion (SI for details on the fit). Mean recoil radius for both MDCK WT and E-cad KO cysts interaction, (solid lines) and corresponding fit for the recoil model (dashed lines). Fit is performed on the mean of (purple, N_WT_=5,n_WT_=9) for WT doublets and (orange, N_KO_=4, n_KO_=23) for E-cad KO doublets. The schematics insert illustrates the baseline stretching state of the monolayer and the associated tension *σ*_base_. Right: Surface velocity of the fusion hole fitted on each early recoil dynamics for WT and E-cad KO doublet fusion (N_WT_=5,n_WT_=9, N_KO_=4, n_KO_=23).

Cyst coalescence is driven by cell-cell adhesion and opposed by viscous dissipation in the deforming monolayer, akin to the coalescence of viscous soap bubbles (Fig. 4B) (29). Cell proliferation plays no significant role, as it would have led to dynamics inconsistent with the measured neck evolution during coalescence (Fig. S7B). In the short-time regime, the neck radius evolves as *a*(*t*) ≈ (*W*_cc_*/η*) *t*, in agreement with experimental data (Fig. 2E). A more general form of *a*(*t*), derived from energy balance considerations (Materials and Methods and SI), captures the full coalescence dynamics (Fig. 4C). The model predicts complete coalescence; however, in experiments, we also observed incomplete coalescence associated to the deposition of matrix (Fig. S3, and SI). This behavior is recapitulated in the model by a gradual reduction in adhesion energy due to extracellular matrix secretion (Fig. S7C). Using the measured rheological parameters, the model predicts coalescence speeds of 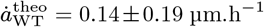 and 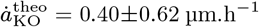, which fall within the experimental range. Fitting the theoretical curve to the neck radius data yields a characteristic time *τ*_v_ = 90 h (Fig. 4C). A scaling argument, *τ*_v_ = *R*_0_*η/W*_cc_ with *R*_0_ the typical radius of the cyst, gives an order of magnitude estimate *τ*_v_ ≈ 100 h, supporting the consistency of the model with measured rheological parameters.

This model based on simple ingredients also provides a rationalization of the proportion of phenotypes observed for WT and E-cad KO cysts (Fig. 1F and Fig. 2B). We describe fusion nucleation as a stochastic process, where active noise triggers the formation of a hole at the interface. The fraction of completely coalesced and fused cyst pairs increases over time with a characteristic rate Γ related to the cyst monolayer tension, here *σ*_asp_ as cells are stretched prior to opening (see Materials and Methods and SI). In Figure 4D, the nucleation rate Γ is fitted to the experimental data, and the model shows excellent agreement, with Γ^KO^ *>* Γ^WT^. Overall reduction in adhesion is translated into a decrease in tension which facilitates fusion.

After nucleation of a hole, lumen fusion proceeds through viscous recoil of the monolayer, resembling the fusion of two low-viscosity bubbles (lumens) within a highly viscous medium (the monolayer) (see schematics Fig. 4E). In the short-time regime, the hole radius increases as *r* = (*V*_s_*/π*)^1*/*2^*t*^1*/*2^ (30), where *V*_s_ represents the velocity of increase of the hole surface, linked to geometrical and mechanical parameters that we measured (see Materials and Methods). The model predicts retraction without rim formation, in agreement with experimental profiles. Figure 4F shows that both WT and E-cad KO cyst data follow the predicted 1*/*2-power law at short times. The best-fit surface velocities *V*_s,WT_ = 72 µm^2^/h and *V*_s,KO_ = 106 µm^2^/h are of the same order of magnitude as the rheology-based estimate (*V*_s_ ≈ 300 µm^2^/h). At longer times, as monolayer tension relaxes, the late recoil velocity matches the characteristic late-time velocity measured in laser cutting experiments (*σ*_cut_*/η* ≈ 0.3 µm/h), consistent with the scaling argument for the rheology-based interpretation.

## Discussion

We report the coalescence and fusion of cyst doublets and explain their dynamics based on the mechanical properties of their cell monolayers. We show that the coalescence velocity is set by the ratio of adhesion energy to viscosity, as is the late fusion velocity, whereas fusion nucleation and early fusion dynamics are influenced by the elastic stretching of the monolayer. Our new doublet assay has been instrumental in studying cyst interaction dynamics, providing spatial and temporal control that enables robust statistics and reliable measurements over more than a week. Our results suggest that rheological parameters capture the dynamics of interaction of MDCK cyst doublets for both WT and E-cadherin KO cysts.

The fact that fusion was facilitated in the case of MDCK E-cad KO cysts can be intuitively understood as weaker junctions allowing enhanced cell-cell remodelling. In contrast, it is surprising that changes in expression of cadherin did not lead to differences in coalescence and interface retraction velocities. Micropipette aspiration experiments revealed differences in monolayer tension and viscosity between the cell lines, as confirmed by the asymmetric coalescence interaction between a WT cyst and a E-cad KO cyst (Fig. S8, Movie S6). However, their ratio remained constant. Previous studies on compact cell aggregates reached similar conclusions (24), suggesting that 3D tissues maintain constant velocities and mechanical parameters despite changes in adhesion levels. The idea of a rheological homeostasis - like tension homeostasis - suggests that cells may adapt changes in adhesion levels to maintain constant speeds for their interactions. This interplay between stress generation in the cortex and adhesion between cells could serve as a guideline for understanding the regulation of tissue dynamics. Our report characterises the physics of cysts through rheological experiments combined with theory, offering new insights into their dynamics. Unlike aggregates, which are compact spheroids of cells without lumens and whose bulk properties have been extensively characterised (20, 24, 25, 31– 33), cysts exhibit a distinct architecture. In terms of soft matter analogies, a compact spheroid can be viewed as a *living drop* (34), whereas a cyst is more appropriately described as a *living bubble*. The physical framework we propose predicts a distinct time evolution of the neck length during coalescence compared to previous studies on cellular aggregates. Earlier aggregate models, using simple liquid or viscoelastic descriptions, predicted short-time dynamics of the neck radius scaling as *a* ∼ *t*^1*/*2^ (4, 23, 35) or as *a* ∼ *t*^1*/*3^ (24). By accounting for the specific geometry of cysts as a cell shell and its associated viscous dissipation volume, our model predicts a different scaling, *a* ∼ *t* (Fig. 2E), with a speed set by the ratio of adhesion energy to viscosity, highlighting a different physical mechanism of viscous dissipation in cyst interactions.

Despite key physical differences between different systems, our rheological measurements align with earlier measurements for different 3D tissues. We retrieve the orders of magnitude of monolayer tension obtained with atomic force microscopy for MDCK cysts, yielding values of 4 mN.m^−1^ (36). Similar monolayer tension values were reported for other cyst types, such as MCF10 cysts (10 mN.m^−1^ (37)), and even intestinal organoids (3-6 mN.m^−1^, with differences between crypts and villi (38)). Comparable tensions have also been measured in compact cell spheroids, with values around 6 mN.m^−1^ (25). Our viscosity order of magnitude is consistent with measurements reported for other hollow structures *in vivo*, such as hydra (2.4.10^6^ Pa.s (39)). We measured a decreased viscosity in MDCK E-cad KO cyst compared to WT cyst, which is consistent with the reported trend for decreased cellular adhesion in embryonic tissue and 2D monolayer (12, 40). These comparisons support the reliability of our measurements and suggest conserved material properties across diverse cell types and systems. Conservations in compositions and stoichiometry of interacting cortical layers across species go along the same concept.

The apparent differences in monolayer tension measured in pipette aspiration and laser cutting experiments can be explained by the degree of monolayer stretching in each case (Fig. 4A). In our rheological characterization, we probed three distinct states: a stretched state, corresponding to nucleation during lumen fusion; a baseline state, corresponding to early fusion dynamics; and a relaxed state, corresponding to late fusion dynamics and coalescence. Importantly, these three degrees of stretching capture the range of mechanical conditions relevant to cyst interactions. We note that the ratio between WT and E-cad KO speeds deduced from lumen cutting experiments differs from that obtained by micropipette aspiration. This deviation likely reflects the larger uncertainty of laser cutting experiments and their sensitivity to cut geometry during drainage, assumed constant in our analysis. Nonetheless, the late-time tension obtained from laser cutting provides the correct order of magnitude for interaction speeds, as predicted by the theoretical model.

While we could have expected the extracellular matrix or hydraulic effects to influence interaction dynamics, our experiments suggest otherwise. Reducing the percentage of Matrigel ruled out a dominant role for the extracellular matrix in setting interaction velocities. Similarly, luminal volume changes are negligible compared to coalescence and fusion monolayer dynamics, indicating that hydraulics plays only a minor role. We also considered an alternative model of coalescence driven by lumen inflation, but its predictions were incompatible with the observed dynamics. Interestingly, the observation of 8-shaped and hourglass doublets reveals an extracellular shell synthesized by the cells that physically constrains cyst shape. Moreover, the probability of achieving cyst fusion appears to be related to the time required to reach full coalescence, suggesting that the rates of matrix synthesis and degradation by the cells may contribute to setting phenotypes. While cell monolayer mechanics appear dominant, future studies should track and quantify matrix dynamics to assess their role, and explore systems with defective epithelial sealing (i.e., permeable to ions) or low lumen occupancy, to further probe the interplay between hydraulic pressure, matrix remodeling, and cyst mechanics.

The extrapolation of our results to *in vivo* systems is appealing but requires caution, as mesenchyme, multiple cellular layers, and matrix secretion or degradation may lead to distinct shape dynamics. Still, the coalescence and fusion stages we observed closely resemble phenomena reported both *in vitro* and *in vivo*: in intestinal organoids, tubes form by fusion of individual cysts with a large lumen volume increase compared to MDCK cysts (41); during angiogenesis, two dorsal aorta merge into the descending aorta, illustrating interactions between tubes as well as cysts (4, 8). Altogether, these similarities suggest that, with minor adaptations, our theoretical approach could help unify these diverse processes within a common physical framework. Extending this work to multilayered systems, such as cysts surrounded by mesenchymal cells or ectodermal–endodermal structures (39), could link the physical behavior of each layer to its active or passive mechanical properties. Reductionist setups like the doublet assay described here may thus serve as valuable tools to test and generalize this mechanical framework, with promising perspectives for synthetic organ design and tissue regeneration (10, 42).

## Materials and Methods

### Cell lines

MDCK type II cells (referred to as MDCK) are cultured on cell culture Petri dishes in MEM with Gibco Minimum Essential Medium (MEM/GlutaMAX) + 5% Fetal Calf Serum (FCS) + 1% Non Essential Amino Acid (100x) + 1mM Sodium Pyruvate + 1% Penicillin-Streptomycin (100x) and maintained at 37°C in 5% CO_2_ incubator with 85% humidity. They are passaged every 2 to 3 days and kept under 80% confluency. MDCK ZO1-mNeonGreen, H2B-mCherry, MDCK E-cadherin-GFP/Podocalyxin-mScarlett/Halo-CAAX (43) that are the reference cell lines for the WT comparison were obtained from the Honigmann lab. For junctional knock-out cell line, we use MDCK E-cadherin KO with no fluorescent marker, from the Ladoux-M‘ege lab (18) to prepare a new stable cell line expressing H2B-mNeonGreen.

### Cell seeding and 3D culture

We used two protocols to obtain cysts, one using cavities and the other based on the culture of cysts between two layers of Matrigel. Before seeding cells in cavities, cavities are plasma activated and incubated with 1ml of laminin (Sigma 11243217001) at a concentration of 5 µg/ml. After detaching cells with trypsin and suspending them in culture medium, cavities are placed in 50ml tube with adaptor to enable centrifugation, with cell in medium on top (17). Cavities and cells are centrifuged 3 times at 300g for 3 minutes. After centrifugation, cavities are washed gently to remove excess cells and 20 µl of Matrigel (Corning, 356231) are placed on top of the cavities and left to polymerize for 10 minutes at 37°C. Culture medium is added to samples. To prepare the sample between two layers of Matrigel (43), glass coverslips are plasma activated. 4 µl of Matrigel are spread as a thin layer on the activated coverslip and left to polymerize for 7 minutes at 37°C. 2000 cells in 4 µl of medium are put on the thin layer of Matrigel and left to adhere for 2 minutes. Remaining culture medium is gently removed and 20 µl of Matrigel is added on top and let to polymerize for 10 minutes at 37°C. Culture medium is added to samples. To test the potential role of Matrigel, we repeated these experiments with 25% Matrigel. We first grew MDCK cysts in 100% Matrigel for 3 days. We then used soft dissociation medium (Stemcell Technologies 100-1077) to remove the Matrigel. To obtain only cysts, medium containing cysts and dissociated Matrigel are centrifuged and we removed supernatant. We then incubated cysts in 20 µl 25% Matrigel diluted in culture medium and let it polymerize for 12 minutes at 37°C. Culture medium is added to samples.

### Microfabrication of cavities

In order to control the growing and interaction conditions of MDCK cysts, we prepared microfabricated cavities with a chessboard design of 6 by 6 large squares containing either cavities doublets for interaction (Fig. 1A, top right) or equally spaced cavities (Fig. 1.A, top left). Briefly, we designed a mask with AutoCAD with large numbers of doublets of cavities with a diameter of 17 µm so that single cells can be trapped. They are separated by 15 µm of inter-distance which lead to interaction of cysts at day 3-4 after seeding of single cells in cavities. A master silicon wafer is prepared using the designed mask. Then performing soft lithography with Polydimethyl-siloxane (PDMS) polymer (Sylgard 184 Silicon elastomer kit), we produce pillars that are used as a transfer mould to make cavities with glass bottom (16, 17).

### Micropipette aspiration

Micropipettes are pulled using the CO_2_ laser based P-2000 Sutter Instrument pipette puller from glass capillaries (Harvard apparatus 30-003) and then cut to the right diameter and bend using a Narishige microforge (Narishige MF-900). Pipettes are passivated using Poly(L-lysine)-graft-poly(ethylene glycol) (PLL(20)-g[3.5]-PEG(2) SuSo) for at least 1 hour at 4°C, and then rinsed with phosphate-buffered saline (PBS). Pressure measurements are performed using the Fluigent Flow EZ −69mbar pressure controller (Fluigent LU-FEZ-N069). Zero pressure is reached when a small difference in pressure, few Pa, is needed to invert the flow at the entrance of the pipette (as described in (26)).

A viscoelastic liquid model of the cell monolayer, treated as a Maxwell liquid with a surface tension, yields the following formulae for the cyst monolayer tension (*σ*_asp_) and viscosity (*η*):

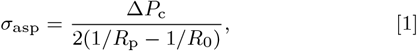

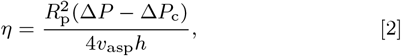

where Δ*P* is the applied pressure difference, Δ*P*_c_ is the critical pressure difference required to make the cyst flow into the pipette, *R*_p_ is the micropipette radius, *R*_0_ is the initial cyst radius, *h* is the monolayer thickness, and *v*_asp_ is the speed of advancement of the aspirated tongue, whose long-time behaviour is fitted by a straight line of slope *v*_asp_. Details of the derivation of these equations are presented in the SI.

### Laser cutting

Cysts were grown until day 4 after seeding in Matrigel. Laser cutting was performed using the UV355nm laser NSV-05P-100 with a power of 5W. Lines for laser cutting ROI were designed such that their lengths were 3 times the height of the monolayer. Laser cutting was done using 10 scanning repetitions and a thickness around the ROI line of 25 in the Modular interface. From the initial outflow *Q* measured from the early volume change after cutting, we compute the hydrostatic pressure difference 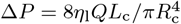 by modelling the outflow as that of a viscous fluid with viscosity *η*_l_, using the Hagen-Poiseuille law for a channel of radius *R*_c_ and length *L*_c_ (28). The baseline monolayer tension is then computed from the Young-Laplace law as *σ*_0_ = *R*_lumen_Δ*P/*2. Similarly, the late-time outflow provides an estimate of the relaxed monolayer tension, *σ*_cut_. We used a luminal fluid viscosity of *η*_l_ = 0.0051 Pa.s, as previously determined (28).

### Microscopy

Mechanical measurements with micropipettes are performed on Olympus wide field microscope with objective 20x air 0.4NA. Samples are imaged in CO_2_ buffered medium Leibovitz’s L-15 supplemented with 5% FCS. Timelapse experiments are performed on the Leica Spinning disk microscope with objective 63x glycerol 1.2 NA. Samples are imaged in passage medium with CO_2_ incubation. We use the 488nm and 561nm laser wavelengths. Cysts were imaged and tracked over days using the Evolus cell culture microscope. Laser cutting was performed on Nikon TiE using and HC PL APO 100x 1.2NA oil objective.

### Data analysis and statistics

Images were analysed using ImageJ. Geometrical measurements were performed on the central cut of cysts doublets. Cell counting was done using the DM3D plug-in and manually counting nuclei on H2B fluorescence channel in 3D. All plots, fit and data analysis were done using Python. Each experiment had at least 3 biological repeats. We used Mann-Whitney test for p-value significance tests. The values in the text are presented as mean ± std.

### Immuno-staining

We followed classical protocols for 3D immunostaining (16, 43). Fixation of the sample was done with 4% paraformaldehyde (PFA) for 10 minutes. Permeabilization was done with Triton x-100 0.5% in PBS at room temperature for 15 minutes. Blocking was performed with normal goat serum at 4°C for 1 day. We used primary antibodies for laminin 111 at 1:50 (L9393 Sigma) incubated for 2 days at 4°C and the secondary antibody (Goat-Anti-Rabbit AlexaFluor 647, Invitrogen A-21245) was used at 1:400 incubated for 2h at room temperature.

### Coalescence model

Cyst coalescence is driven by the rate of energy gain from forming the interface, 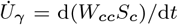, where *W*_cc_ is the cell-cell adhesion energy per unit interface and *S*_c_ = *πa*^2^ is the contact area, with *a* = *R* sin *θ* the neck radius (see Fig. 4B). Viscous dissipation 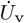 opposes coalescence and is concentrated at the edges of the contact region (cells marked in blue in Fig. 4B). We thus write 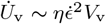,where *η* is the dynamic viscosity, 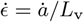 the deformation rate, *L*_v_ the characteristic dissipation length (of the order of the cell size), and *V*_v_ = 2*πaL*^2^ the dissipation volume. Balancing energy gain and dissipation 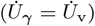 yields the evolution of *a*(*t*). Since cyst growth and lumen hydraulic loading do not play a dominant role in cyst coalescence (see SI), we assume lumen volume conservation over the time scale for adhesion-driven coalescence, which provides the evolution equation for the lumen radius, *R*(*t*). Together with reasonable geometric simplifications (see SI), this leads to the approximate differential equation:

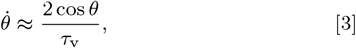

where *τ*_v_ = *ηR*_0_*/W*_cc_ is the characteristic viscoelastic time and *R*_0_ is the initial radius. The solution,

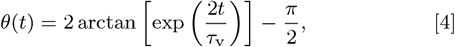

describes the experimental data well (Fig. 4C). At short times (*t* ≪ *τ*_v_) Eq. 4 simplifies to *θ* = 2*t/τ*_v_, corresponding to *a* = 2*R*_0_*t/τ*_v_, predicting linear coalescence consistent with experiments (Fig. 2E).

### Fusion nucleation model

Hole nucleation in the monolayer is modeled as a noise-driven process (44, 45). The nucleation rate is:

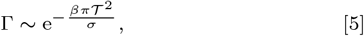

where *β*^−1^ quantifies the magnitude of the active noise, 𝒯 is the line tension opposing hole formation, and *σ* the monolayer tension. Fusion of a population of cyst doublets is treated as a *linear death process* (46), where the number of non-fused doublets follows a binomial distribution. At long enough time so that coalescence is assumed completed, the fraction of fused doublets evolves as:

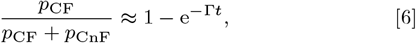

with *p*_CF_ and *p*_CnF_ denoting the fractions of fused and not fused doublets, respectively. This relation is fitted to experimental data to extract the nucleation rate Γ. Fitted Γ values are consistent with rheological measurements, with *σ* = *σ*_asp_ since hole nucleation occurs in a stretched region of the monolayer, where cell stretching is comparable to that observed during micropipette aspiration, *L* ≈ 1.5*L*_base_ (Fig. S5B, Fig. S6 and SI for details).

### Lumen fusion model

Lumen fusion is modeled as the viscous recoil of a cell monolayer, similar to the fusion of two low-viscosity bubbles in a highly viscous fluid (30) In the Stokes limit, the fusion pore radius evolves as *r*(*t*) = (*V*_s_*/π*)^1*/*2^*t*^1*/*2^, where *V*_s_ = *Cπ*(*Rσ/η*), with *R* the cyst radius, *σ* and *η* the tension and viscosity of the monolayer, and *C* = 0.7935 in the inertialess limit. The surface velocity *V*_s_ was thus estimated from rheological measurements, using *σ* = *σ*_base_ as an approximation for the average monolayer tension during recoil.

## ACKNOWLEDGMENTS

We thank R. Berthoz, T. Guyomar and the Riveline group for help and discussions, and the Imaging Platform of IGBMC. We thank B. Ladoux and A. Honigmann for providing some of the MDCK cell lines. D.R. acknowledges the Interdisciplinary Thematic Institute IMCBio, part of the ITI 2021-2028 program of the University of Strasbourg, CNRS, and Inserm, which was supported by IdEx Unistra (ANR-10-IDEX-0002), and by SFRI-STRAT’US project (ANR 20-SFRI-0012) and EUR IMCBio (ANR-17-EURE-0023) under the framework of the French Investments for the Future Program. D.R. is supported by Research Grants from USIAS, Labex, ANR, SNF Sinergia grant CRSII5 183550 and the Human Frontier Science Program RGP0050/2018.

