## Supplementary material for "Interaction dynamics between epithelial cysts captured by tissue rheology": Suppl. Information

#### This PDF file includes:

Supporting text
Figs. S1 to S9
Legends for Movies S1 to S6
SI References

#### Other supporting materials for this manuscript include the following:

Movies S1 to S6

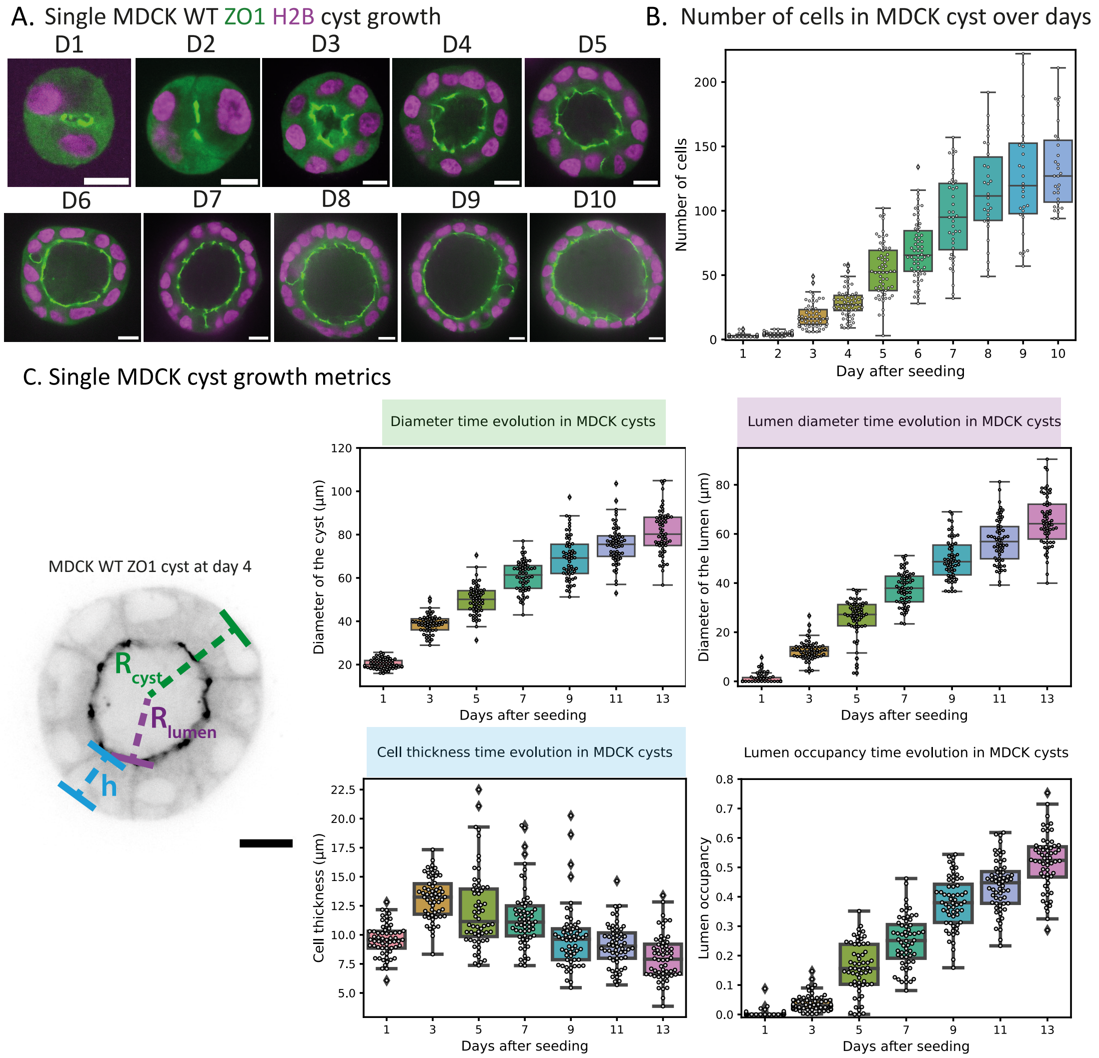


**Fig. S1.** Characterisation of single MDCK cysts growth. A) MDCK cyst ZO1-mNeonGreen (green) H2B-mCherry (magenta) imaged each day for 10 days, scale bars 10 µm. B) Cell number over days of growth of MDCK cysts (N=3, n=30-60). C) Left: Geometrical description of single epithelial cyst with the radius of the cyst $R_{\mathrm{cyst}}$, the radius of the lumen $R_{\mathrm{lumen}}$ and the cell height $h = R_{\mathrm{cyst}} - R_{\mathrm{lumen}}$. MDCK cyst ZO1-mNeonGreen aged day 4 after seeding presented in inverted contrast, scale bar 10 µm. Center top: cyst diameter over days. Right top: lumen diameter over days. Center bottom: cell height over days, Right bottom: lumen occupancy $R^2_{\mathrm{lumen}}/R^2_{\mathrm{cyst}}$ over days (N=3 and n=60).

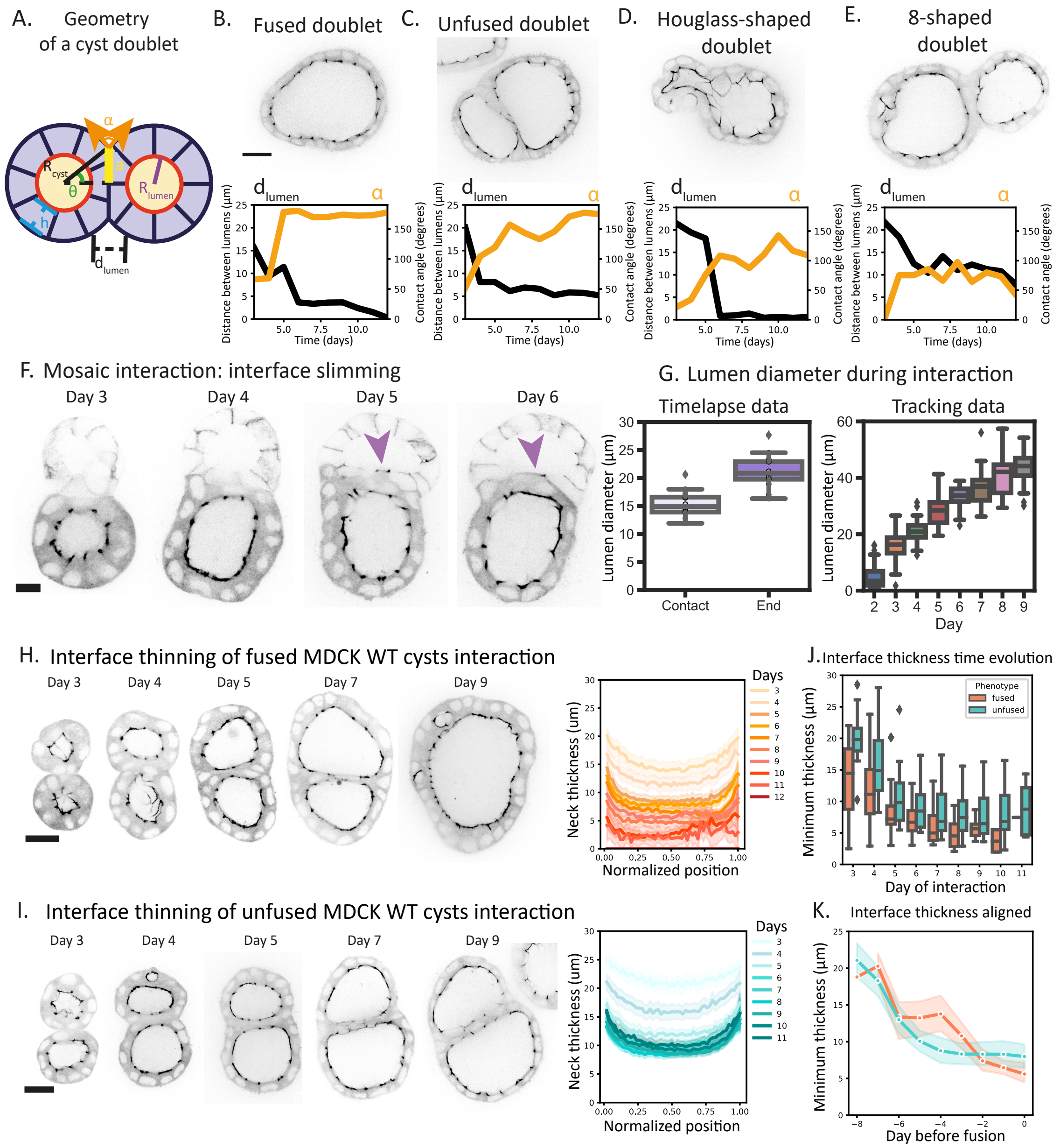


**Fig. S2.** Interaction phenotype geometrical signatures and interface evolution during the interaction between cysts. A) Schematics of interacting MDCK cysts. We define the geometry of a doublet of cysts is described by the radius of each cyst $R_{cyst}$, the radii of the lumens $R_{lumen}$, the radius of the interaction area a, the contact angle $\alpha$, the thickness of the interface between two lumen $d_{lumen}$ and h the thickness of the monolayer of cell. Blue regions indicate the most deforming cells during the interaction. B) Geometrical characterization of fused doublet phenotype. Top: fused doublet phenotype of interaction between MDCK WT cysts at day 11. Bottom: Distance between lumens and contact angle. C) Geometrical characterization of unfused doublet phenotype. Top: unfused doublet phenotype of interaction between MDCK WT cysts at day 11. Bottom: Distance between lumens and contact angle. D) Geometrical characterization of hourglass-shaped doublet phenotype. Top: hourglass-shaped doublet phenotype of interaction between MDCK WT cysts at day 11. Bottom: Distance between lumens and contact angle. E) Geometrical characterization of 8-shaped doublet phenotype. Top: 8-shaped doublet phenotype of interaction between MDCK WT cysts at day 11. Bottom: Distance between lumens and contact angle. F) Mosaic interaction of two MDCK WT cysts expressing E-cadherin-mNeonGreen (top cyst) and ZO1-mNeonGreen (bottom cyst). The purple arrows indicate the cell from the bottom cyst located at the interface that intercalates with the top cyst cells. From day 4 to day 6, the cysts doublet goes from two layers of cells at the interface to a single one. This transition is highlighted by the ZO1 signal (located apically) present on both sides of the intercalating cell. Scale bar 10 µm. G) Lumen diameter in um during interaction of MDCK WT cysts on both timelapse (interaction starts at day 2.5 and finishes around day 4, N=4,n=14) and tracking data (N=3,n=24). H) Left: Interaction of two MDCK WT cysts expressing ZO1-mNeonGreen undergoing thinning of the interface and fusion of lumens. Scale bar 20 µm. Right: Neck thickness in µm along the normalized interacting cysts interface with time for fused lumens interaction phenotypes. Lighter colours indicate earlier days of interaction. Thickness along the interface is plotted as mean ± 95% confidence interval (N=5, n=33). I) Left: Interaction of two MDCK WT cysts expressing ZO1-mNeonGreen undergoing incomplete thinning of the interface and no fusion of lumens at day 11. Scale bar 20 µm. Right: Neck thickness in µm along the normalized interacting cysts interface with time for unfused lumens interaction phenotypes. Lighter colours indicate earlier days of interaction. Thickness along the interface is plotted as mean ± 95% confidence interval (N=3, n=21). J) Interface minimum thickness time evolution during the interaction for both fused (orange) and unfused (blue) phenotypes at day 11. K) Interface minimum thickness time evolution during the interaction for both fused (orange) and unfused (blue) phenotypes aligned on the day of fusion and plotted as mean ± 95% confidence interval.

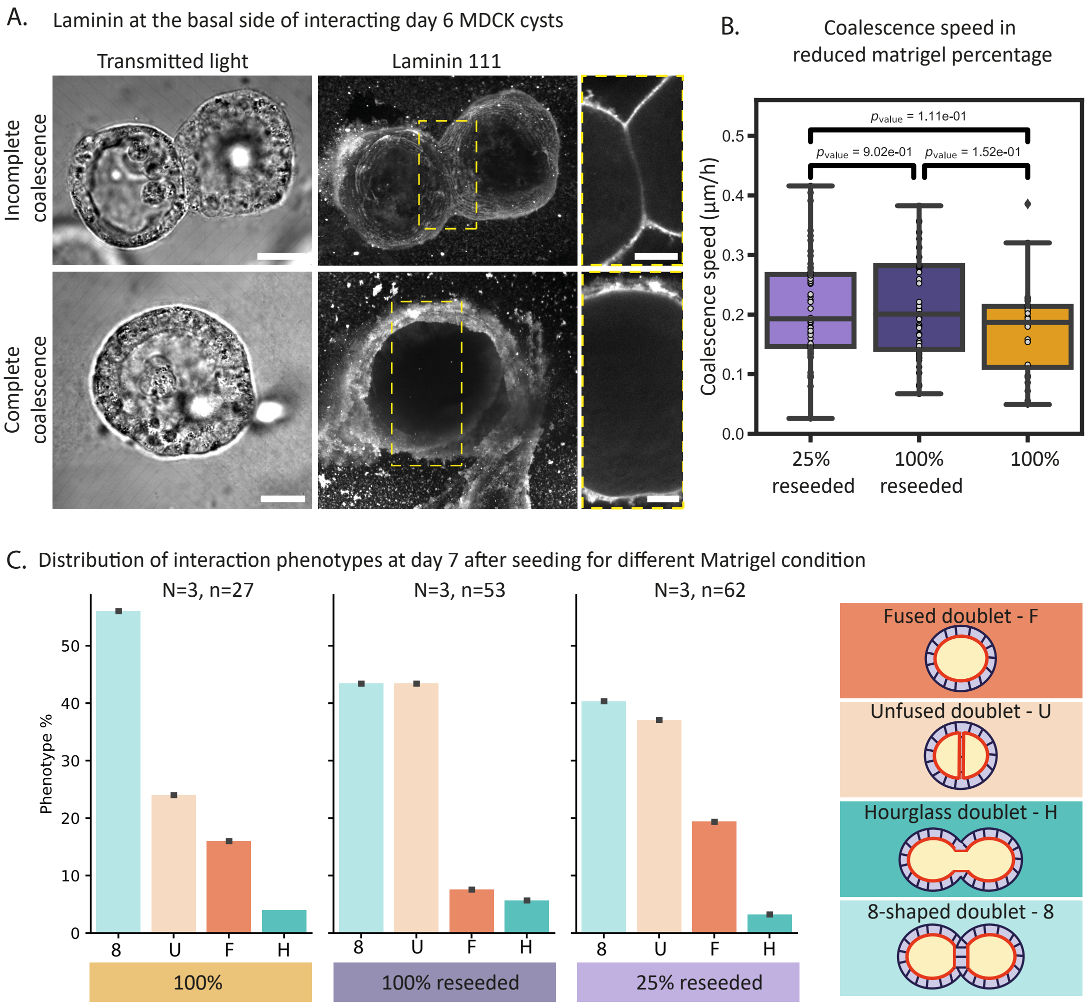


**Fig. S3.** Coalescence speed is not affected by the Matrigel percentage. A) Location of laminin in interacting MDCK cyst doublets either incompletely coalesced (top) or completely coalesced (bottom). The sample is fixed and stained for laminin at day 6 after seeding. The laminin images are maximal intensity projections that highlight the presence of basal laminin. The cropped images correspond to the central slice of the yellow dashed rectangle on the laminin image. scale bars 20 μm for non cropped images and 10 μm for cropped images. B) Coalescence speeds measured for MDCK WT interaction starting at day 3 after seeding in 100% Matrigel. Cysts are dissociated from the matrix and reseeded in either 100% or 25% of Matrigel diluted in culture medium and compared to normal interaction of cysts in 100% Matrigel (N=3 for each and $n_{100\%} = 27$, $n_{100\% reseed} = 53$, $n_{25\% reseed} = 62$). C) Distribution of interaction phenotypes at day 7 after seeding for the different Matrigel culture conditions presented in B). F stands for fused doublets, U for unfused doublet, H for hourglass doublets and 8 for 8-shaped doublet (see Fig 1G).

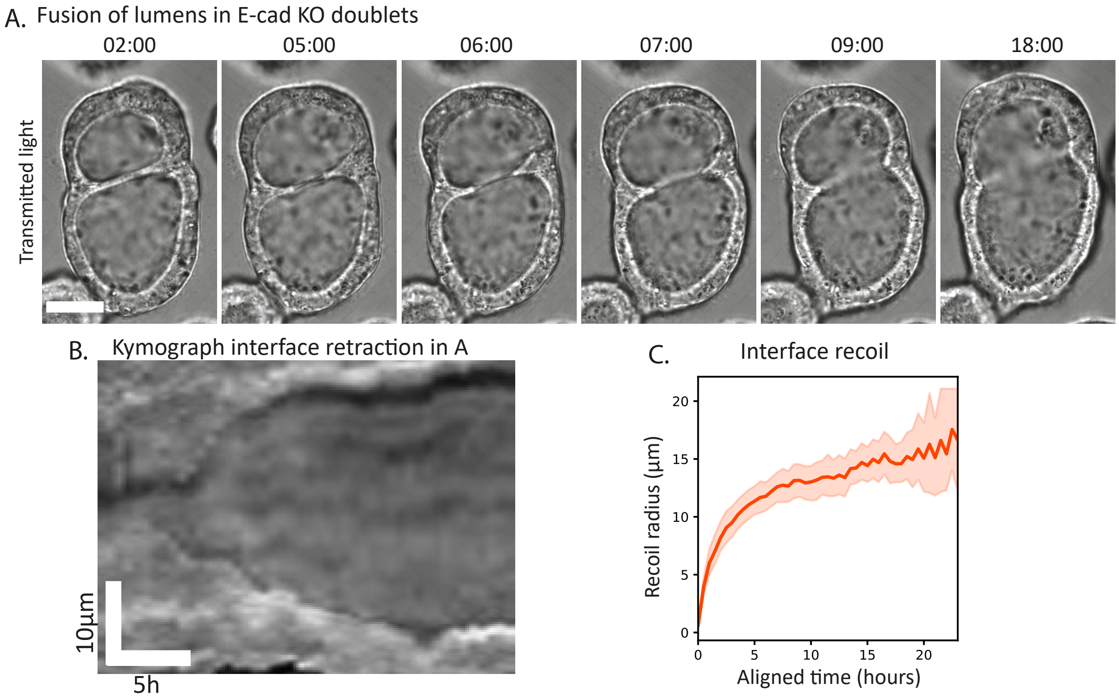


**Fig. S4.** Fusion of lumens in MDCK E-cad KO cysts doublets. A) Fusion of lumens MDCK E-cad KO cysts doublets aged day 4 after seeding between two layers of Matrigel. Time in hh:mm, scale bar 20 µm. B) Kymograph of the lumen fusion presented in A. The darker lines correspond to the interface recoil. Time reads along the horizontal axis and position along the vertical axis. C) Interface recoil length in µm over time during lumen fusion of MDCK E-cad KO cysts interaction (N=4, n=23).

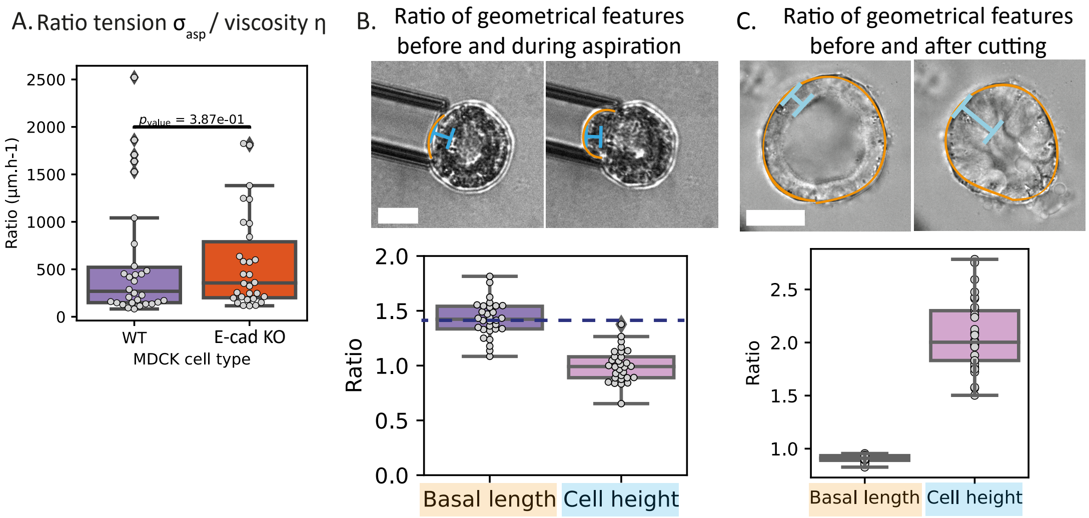


**Fig. S5.** Stretching of monolayers in the different rheological experiments. A) Ratio between the effective monolayer tension and the viscosity in µm.h-1 measured through micropipette aspiration (N=3, n=30). B) Top: Basal stretching of MDCK cysts aged day 4 after seeding during micropipette aspiration. The orange line corresponds to the basal length measured before aspiration and during aspiration used to compute stretching of the monolayer in B. The blue lines correspond to the cell height measured before aspiration and during aspiration. Scale bar 20 µm. Movie S3. Bottom: Ratio of basal lengths before/during aspiration and cell height before/during aspiration. The dashed line indicates a ratio value of $1.4 \approx \sqrt{2}$ corresponding to a basal stretch doubling the basal area (N=3, n=30). C) Top: Laser piercing of MDCK WT cysts aged day 4 after seeding (left: before piercing, right: after piercing). The blue line corresponds to the cell height and the orange outline corresponds to the basal length. Scale bar 20 µm. Movie S5. Bottom: Ratio of basal lengths before/after piercing and cell height before/during aspiration (N=3, n=28).

 

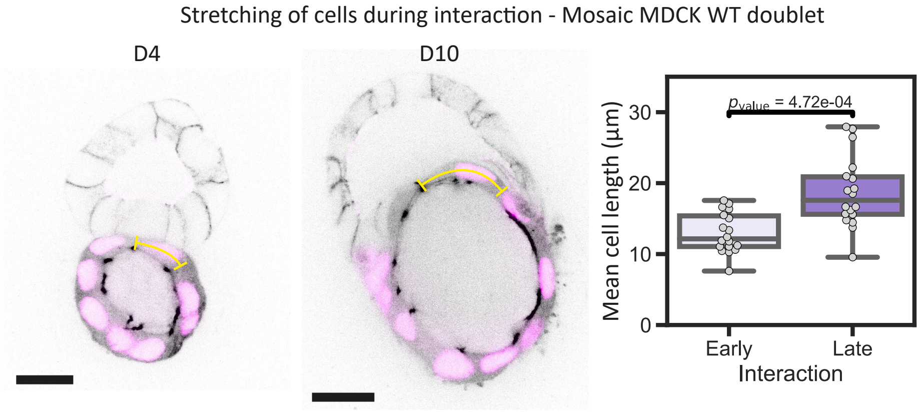


**Fig. S6.** Stretching of cells in the interaction neck. Left and centre: mosaic interaction of WT cysts doublets highlighting the stretching of cells in the neck region. The top cyst expresses fluorescently E-cadherin and podocalyxin and the bottom one ZO1 and H2B. Right: Quantification of mean cell elongation during the interaction, early and late (N=3 and n=18). Scale bars 20 µm.

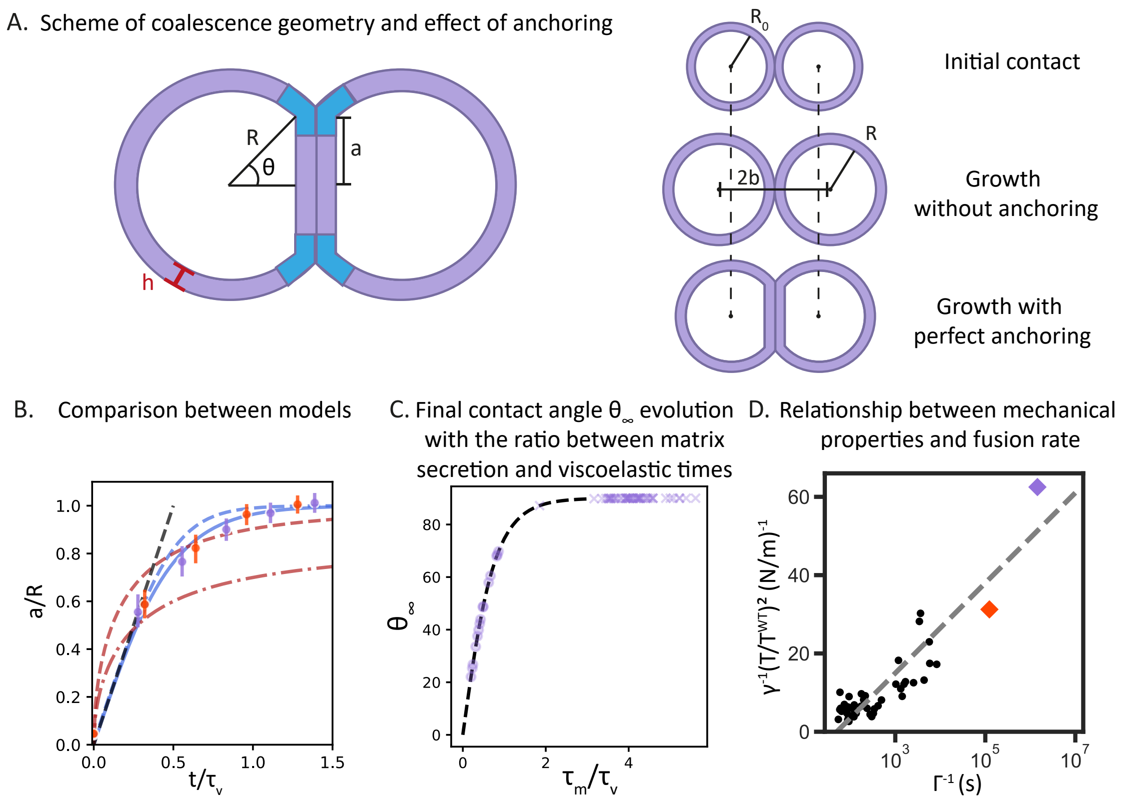


**Fig. S7.** Physical modeling of cysts interaction (see SI theory part). A) Left: Schematics of the geometrical description of cyst coalescence. The blue regions indicate the cells that concentrate the viscous dissipation. Right: Geometrical evolution of a growing cyst doublet in the absence or presence of anchoring. B) Comparison between the growth-driven coalescence and adhesion-driven coalescence normalized neck radius, $a/R$. Growth-driven coalescence for $\alpha$ = 0 in red dashed line and $\alpha$ = 0.5 in red dash-dotted line. Predicted adhesion-driven coalescence with dotted line for the exact solution of Eq. 19, full blue line for the approximation of Eq. 21, straight dashed line for the linear approximation at short time. Experimental data for WT and E-cad KO interaction are in purple and orange, respectively. C) Final coalescence angle $\theta_\infty$ as a function of the ratio of matrix secretion to viscoelastic timescales, $\tau_\mathrm{m}/\tau_\mathrm{v}$. Values of $\theta_\infty$ smaller than $\pi/2$ correspond to incomplete coalescence. Experimental data for WT doublets were fitted on the model depending on their final coalescence angle. Circles correspond to incomplete coalescence and crosses to complete coalescence. D) Relationship between mechanical properties and the fusion rate $\Gamma$, including our experiments (WT and E-cad KO) and data from Duque et al. (1). Nucleation theory predicts a linear relation between $\mathcal{T}^2/\gamma$ and log($\Gamma^{-1}$), indicated by the dotted line.

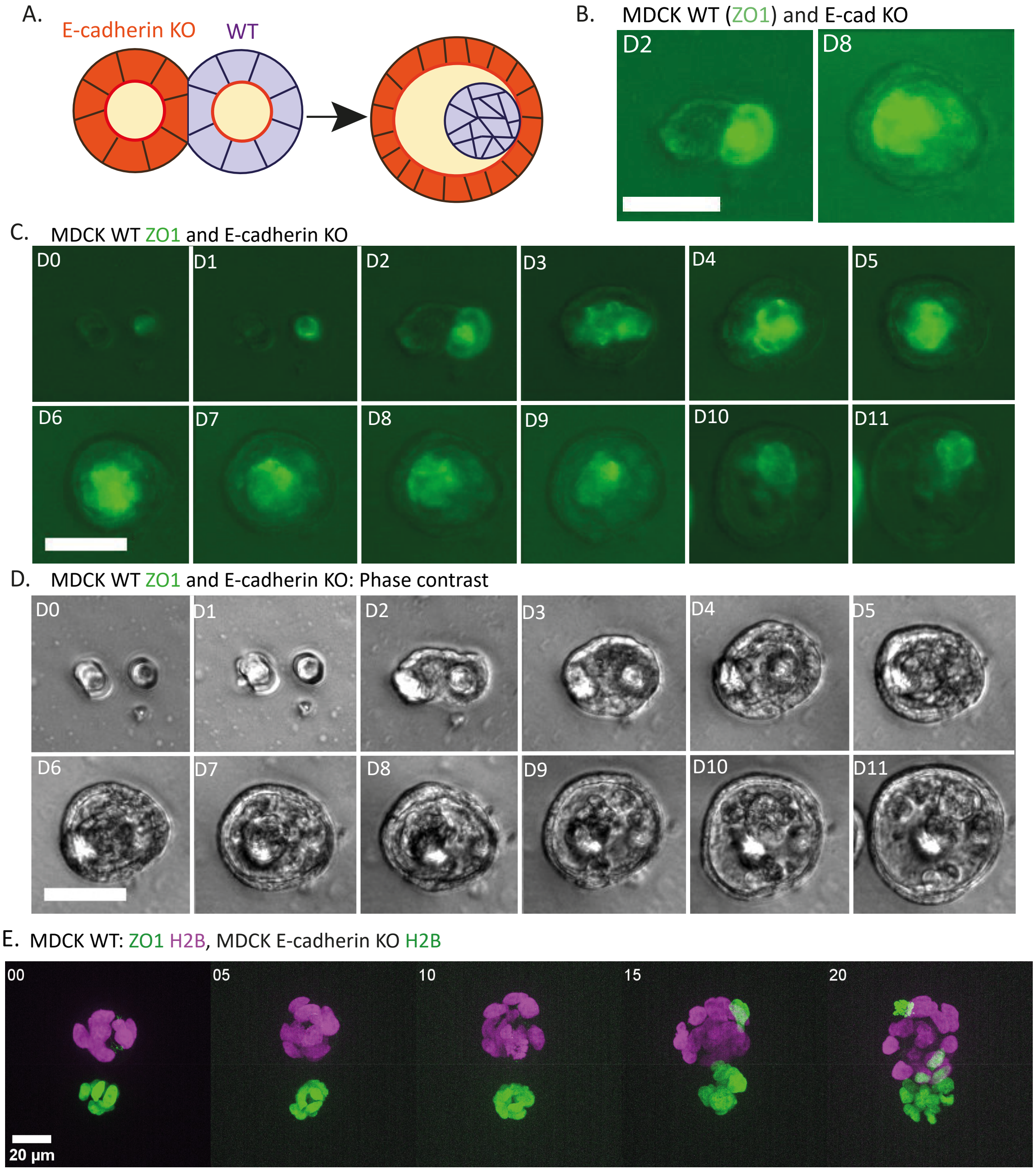


**Fig. S8.** Engulfment of MDCK WT by E-cadherin KO cysts during interaction. A) Schematics representing the engulfment process. MDCK E-cadherin KO cyst is indicated as orange (and has the lowest surface tension), MDCK WT cyst is indicated in purple (and has the highest surface tension). WT engulfed cysts lose the apical-basal polarity. B) Engulfment process on tracking acquisition. Data corresponds to the full process shown in C. Scale bar 50 µm. C) Full engulfment process on tracking acquisition of WT/E-cad KO doublet interaction over 11 days of interaction. Fluorescence channel corresponds to the WT ZO1-mNeonGreen cells. Scale bar 50 µm. D) Full engulfment process on tracking acquisition of WT/E-cad KO doublet interaction over 11 days of interaction. Phase contrast images. Scale bar 50 µm. E) Timelapse interaction process of WT/E-cad KO doublet. An E-cad KO H2B green cell initiates the engulfment process by moving around the WT cyst. Scale bar 20 µm. Movie S6.

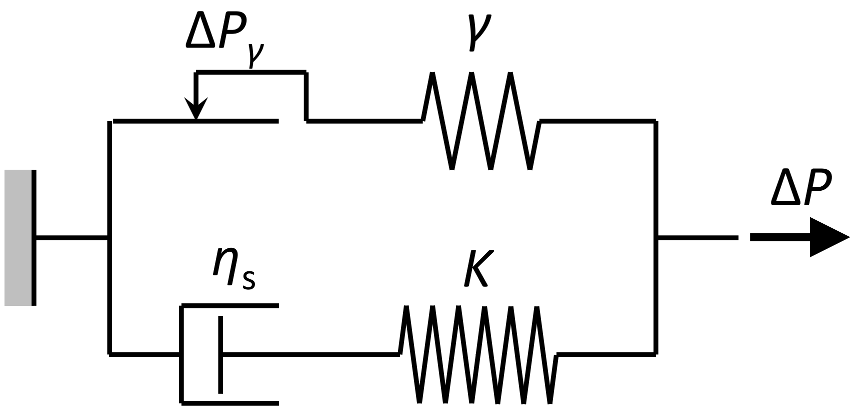


**Fig. S9.** Schematics of the cyst rheological behavior during micropipette aspiration

### Supporting Information Text

### Theoretical models

**Dynamics of cyst aspiration.** To extract rheological parameters from micropipette aspiration experiments, we adapt the formalism proposed by Hochmuth and Berk for membrane tube pulling (2). We conceptualize the aspirated cyst tongue as a cylinder of length $l$ and radius $R_\mathrm{p}$. Assuming local area conservation, continuity requires

$$v_r = -\frac{\dot{l} R_\mathrm{p}}{r}, \quad [1]$$

where $v_r$ is the monolayer velocity in the radial direction $r$, perpendicular to the pipette axis.

We model the monolayer as a 2D Maxwell liquid possessing a surface tension, as schematized in Fig. S9. In the rheological model shown in the figure, the upper branch describes the cyst's surface tension by a combination of a spring of constant $\gamma$ and a Saint-Venant element that slides with constant friction $\Delta P_\gamma$. The lower branch captures the long-time flow of the cyst as a Maxwell liquid of elastic constant $K$, corresponding to the area elastic modulus, and 2D viscosity $\eta_\mathrm{s}$. The total tension in the cyst monolayer is the sum of the surface tension and elastic contributions (3, 4). In a linear areal elasticity approximation,

$$\sigma = \gamma + K\left(\frac{A}{A_0} - 1\right), \quad [2]$$

where $\gamma$ is the surface tension, $K$ the area elastic modulus, and $A/A_0$ is the areal stretch. The minimal aspiration pressure required to induce cyst flow into the micropipette, $\Delta P_\mathrm{c}$, induces an aspirated length $l = R_\mathrm{p}$, corresponding to areal stretch $A/A_0 \approx 2$. Therefore, the effective monolayer tension at the flow threshold is $\sigma_\mathrm{asp} = \gamma + K$. According to Laplace's law, the critical aspiration pressure is then

$$\Delta P_\mathrm{c} = 2\sigma_\mathrm{asp}\left(\frac{1}{R_\mathrm{p}} - \frac{1}{R_0}\right), \quad [3]$$

where $R_0$ is the initial lumen radius. By measuring $\Delta P_\mathrm{c}$, the critical pressure above which a cyst flows into the micropipette, Eq. 3 allows calculating the effective monolayer tension, $\sigma_\mathrm{asp}$.

For $\Delta P > \Delta P_\mathrm{c}$, the cyst flows into the pipette. After an initial transient regime observed in the experiment, the aspiration dynamics are well described by the equation $l(t) = l_0 + v_\mathrm{asp} t$, characteristic of a Maxwell liquid. This behavior is captured by the rheological model in Fig. S9 providing that we set the value $\Delta P_\gamma = 2\gamma(1/R_\mathrm{p} - 1/R_0)$ to remain consistent with the expression of $\Delta P_\mathrm{c}$ deduced in Eq. 3 above. The intercept $l(t=0) = l_0$ corresponds to the elastic radial stress,

$$T_{r,\mathrm{el}} \approx \gamma + K\left(\frac{1\pi R_\mathrm{p} l_0}{\pi R_\mathrm{p}^2} - 1\right). \quad [4]$$

Force balance requires

$$T_{r,\mathrm{el}} 2\pi R_\mathrm{p} = \left(\Delta P - \frac{2\sigma_\mathrm{asp}}{R_0}\right)\pi R_\mathrm{p}^2, \quad [5]$$

which leads to

$$K = \frac{R_\mathrm{p}^2(\Delta P - \Delta P_\mathrm{c})}{4(l_0 - R_\mathrm{p})}. \quad [6]$$

At long time, the cyst flows as a viscous fluid. In this limit, the viscoelastic radial stress can be written as

$$T_r = \sigma_\mathrm{asp} + 2\eta_\mathrm{s}\frac{\partial v_r}{\partial r}, \quad [7]$$

where $\eta_\mathrm{s}$ is the 2D surface viscosity, with units of Pa.m.s. Assuming that radial stress is continuous at the shell turn into the micropipette and neglecting friction with the micropipette walls, force balance reads

$$T_r|_{r=R_\mathrm{p}} 2\pi R_\mathrm{p} = \left(\Delta P - \frac{2\sigma_\mathrm{asp}}{R}\right)\pi R_\mathrm{p}^2, \quad [8]$$

where $\Delta P$ is the aspiration pressure and $R$ is the lumen radius outside the micropipette. Making use of Eqs. 1, 3, 7 and 8, we obtain the relationship

$$\Delta P - \Delta P_\mathrm{c}\left(\frac{1 + R_\mathrm{p}/R}{1 + R_\mathrm{p}/R_0}\right) = \frac{4\eta_\mathrm{s}}{R_\mathrm{p}^2}\dot{l}. \quad [9]$$

By assuming $R \approx R_0$, we obtain the following equation describing the aspiration dynamics:

$$\dot{l} = \frac{R_\mathrm{p}^2}{4\eta_\mathrm{s}}(\Delta P - \Delta P_\mathrm{c}). \quad [10]$$

By measuring the speed of cyst aspiration into the micropipette, $\dot{l} = v_\mathrm{asp}$, Eq. 10 allows calculating the 2D viscosity of the cyst monolayer, $\eta_\mathrm{s}$. The corresponding bulk viscosity is then obtained as $\eta = \eta_\mathrm{s}/h$, where $h$ is the monolayer thickness.

**Growth-driven cyst coalescence.** Here we investigate the hypothesis that coalescence dynamics is determined by cyst growth. As a first step, we must identify the mechanism underlying cyst growth in our setup. An intuitive but naive hypothesis is that cyst growth is driven by cell proliferation at a constant rate. This would imply an exponential growth of the cyst size, inconsistent with experimental observations. A second hypothesis is that cyst growth is set by lumen swelling, a hydraulic mechanism that we examine below.

Lumen swelling results from water flow into the cyst, which is driven by osmotic and hydrostatic pressure differences and is limited by cell permeability. The temporal evolution of the lumen radius $R$ is described by the following equation (4, 5):

$$\frac{\mathrm{d}R}{\mathrm{d}t} = \lambda_{\mathrm{w}}(\Delta\Pi - \Delta P), \quad [11]$$

where $\lambda_{\mathrm{w}}$ is the cell permeability to water (of the order of $10^6$–$10^7$ µm·s$^{-1}$·Pa$^{-1}$ for MDCK cells (4)) and $\Delta P$ and $\Delta\Pi$ are respectively the hydrostatic and osmotic pressure differences between the lumen and the outer medium. Equation 11 can be simplified by the two following considerations. First, for soft materials such as the cyst, we expect $\Delta P \ll \Delta\Pi$ (5). Indeed, we expect $\Delta\Pi \approx 3$–$300$ kPa, whereas $\Delta P \approx 20$–$300$ Pa (4, 6). Second, because cells actively stabilize osmolite lumen concentration fast (over a time scale of minutes (7)) as compared to the time scale of lumen evolution, we can assume that $\Delta\Pi$ remains approximately constant over time. Thus, Eq. 11 simplifies to $\mathrm{d}R/\mathrm{d}t \approx \lambda_w \Delta\Pi$, leading to

$$R = \lambda_{\mathrm{w}}\Delta\Pi(t + t_0), \quad [12]$$

where $t_0$ is an adjustable parameter. This result corresponds to a linear increase of the lumen radius over time, with a characteristic velocity of the order of several micrometers per day, which is consistent with the experimentally observed growth dynamics over 9 days for both isolated and coalescing cysts (Fig. S1C and S2G). In contrast, if cell proliferation were the dominant mechanism for cyst growth, one would expect an exponential cyst size evolution, corresponding to a constant rate of cell proliferation. These results support the hypothesis that cyst growth is primarily governed by the hydraulics of lumen swelling.

We next investigate the effect of hydraulic-driven growth on cyst coalescence. Growth of a cyst doublet coupled with perfect anchoring of the cyst centres, which can be provided by the substrate anchors or by the surrounding Matrigel, will lead to a geometry evolution that resembles coalescence, as illustrated in Fig. S7A. Note however that, in the absence of anchoring, growth will not induce coalescence. Rather, if coalescence occurs, growth without anchoring will slow it down, since it will decrease the ratio $a/R$ (see Fig. S7A for a definition of the geometrical parameters used to describe cyst coalescence).

A purely geometric, growth-induced coalescence can be modeled by writing $b$, the half-distance between the cyst centers (see Fig. S7.A), as:

$$b(t) = R_0 + \alpha(R(t) - R_0), \quad [13]$$

where $R_0$ is the cyst radius at initial contact, $R(t) = R_0 + \lambda_{\mathrm{w}}\Delta\Pi\, t$ is the cyst radius at time $t$ after contact, and the parameter $\alpha$ quantifies the cyst anchoring strength, with $\alpha = 0$ for perfect anchoring and $\alpha = 1$ for no anchoring. Then,

$$\theta(t) = \arccos\frac{b(t)}{R(t)} = \arccos\frac{R_0 + \alpha\lambda_{\mathrm{w}}\Delta\Pi}{R_0 + \lambda_{\mathrm{w}}\Delta\Pi}. \quad [14]$$

Equation 14 is compared to experimental data in Fig. S7B. For perfect anchoring ($\alpha = 0$, dashed red line), Eq. 14 predicts a very fast initial coalescence followed by a significant slowdown, a shape that is inconsistent with experimental observations (symbols in Fig. S7B). The agreement is equally bad if we assume partial anchoring (e.g., $\alpha = 1/2$, dash-dotted red line). The disagreement between predictions and data indicates that hydraulics-driven cyst growth is not the primary mechanism of the observed coalescence dynamics, even if the fast initial growth-induced dynamics may explain the very fast initial coalescence observed at very short times (initial 5 hours, see Fig. 2E in the main text).

**Adhesion-driven cyst coalescence.** Since a model based on hydraulics-driven growth fails to reproduce the measured coalescence dynamics, we next investigate the dynamics of adhesion-driven cyst coalescence. In this scenario, coalescence is driven by cell adhesion between both cysts and opposed by viscous dissipation resulting from cell deformation, akin to the coalescence of viscous soap bubbles (8). Having established that cyst growth does not play a dominant role in cyst coalescence, here we assume that lumen volume remains approximately constant over the relevant time scale for adhesion-driven coalescence (as supported by measurements in Fig. S2F). Lumen volume conservation leads to the relationship (9):

$$R(\theta) = 2^{2/3}(1 + \cos\theta)^{-2/3}(2 - \cos\theta)^{-1/3}R_0, \quad [15]$$

where the geometrical parameters are defined in Fig. S7A, with $R$ the lumen radius and $R_0 = R(t = 0)$ the initial lumen radius. Cyst coalescence is driven by the rate of energy gain due to creation of the interface, $\dot{U}_\gamma$, and resisted by viscous dissipation, $\dot{U}_{\mathrm{v}}$. Let us discuss these two terms separately.

The rate of energy gain due to creation of the interface is

$$\dot{U}_\gamma = \frac{d(W_{cc}S_c)}{dt}, \quad [16]$$

where $W_{\mathrm{cc}} = \gamma - \gamma_{\mathrm{cc}}$ is the energy gained per unit interface, with $\gamma$ the apico-basal cyst surface tension and $\gamma_{\mathrm{cc}}$ the cell-cell tension at each of the two layers that form the interface. We approximate $W_{\mathrm{cc}} \approx \gamma$, assuming $\gamma_{\mathrm{cc}} \ll \gamma$ (10). $S_{\mathrm{c}} = \pi a^2$ is the contact surface, with $a = R\sin\theta$ the radius of the neck (see Fig. S7A). As we will later discuss, we expect the effective adhesion affinity to decrease over time, as adhesion becomes impaired by the secretion of extracellular matrix, which will eventually lead us to consider a time-varying effective $W_{\mathrm{cc}}$.

To compute the rate of viscous dissipation we consider that, at a given coalescence stage, the cells that are deforming the most are those at the edges of the contact region (cells marked in blue in Fig. S7A), and therefore they concentrate viscous dissipation. We thus write

$$\dot{U}_{\mathrm{v}} = A\eta\dot{\epsilon}^2 V_{\mathrm{v}}, \tag{17}$$

where $A = O(1)$ is a numerical coefficient that depends on the details of the viscous flow, $\eta$ is the dynamic viscosity, $\dot{\epsilon} = \dot{a}/L_{\mathrm{v}}$ is the deformation rate, with $L_{\mathrm{v}}$ the width of the dissipation region (of the order of the cell size), and $V_{\mathrm{v}} = 2\pi a L_{\mathrm{v}}^2$ is the volume of the dissipation region.

Cyst coalescence is governed by the equation

$$\dot{U}_\gamma = \dot{U}_{\mathrm{v}}, \tag{18}$$

which leads to the differential equation

$$\dot{\theta} = \frac{W_{\mathrm{cc}}}{\eta R(t)} 2\cos\theta(2-\cos\theta), \tag{19}$$

with $R(t)$ given by Eq. 15. As proposed by Flenner et al. (9), a simpler, approximate solution can be obtained by making $R \approx R_0$ and $2-\cos\theta \approx 1$, leading to

$$\dot{\theta} \approx \frac{2\cos\theta}{\tau}, \tag{20}$$

where $\tau_{\mathrm{v}} = \eta R_0/W_{\mathrm{cc}}$ is the characteristic viscoelastic time. The solution of Eq. 20 is

$$\theta(t) = 2\arctan\left[\exp\left(\frac{2t}{\tau_{\mathrm{v}}}\right)\right] - \frac{\pi}{2}. \tag{21}$$

At short time, $t \ll \tau_{\mathrm{v}}$, Eq. 21 simplifies to $\theta = 2t/\tau_{\mathrm{v}}$, corresponding to $a = 2R_0 t/\tau_{\mathrm{v}}$, i.e., linear coalescence. At long time, Eq. 21 approaches the asymptotic values $\theta \to \pi/2$ and $a \to R$, corresponding to full coalescence. Predictions of this adhesion model are shown in Fig. S7B. The exact solution of Eq. 20 (dotted blue lines) is reasonably represented by the approximate solution of Eq. 21 (solid blue lines), and they both appropriately represent the experimental observations. The figure also shows the predicted short-time linear behavior (highlighted by the dashed black line), which is confirmed experimentally (Fig. 2E in the main text, between 5h and 20h).

**Extracellular matrix secretion may lead to incomplete coalescence.** We account for the effect of ECM secretion on cyst coalescence by an effective, reduced adhesion energy $W_{\mathrm{eff}}$, which we write

$$W_{\mathrm{eff}} = W_{\mathrm{cc}} \exp\left(-t/\tau_{\mathrm{m}}\right), \tag{22}$$

where $\tau_{\mathrm{m}}$ is the characteristic time of extracellular matrix secretion. Introducing this definition into the energy balance equation leads to

$$\theta(t) = 2\arctan\left\{\exp\left[2\frac{\tau_{\mathrm{m}}}{\tau_{\mathrm{v}}}\left(1-\mathrm{e}^{-t/\tau_{\mathrm{m}}}\right)\right]\right\} - \frac{\pi}{2}. \tag{23}$$

As shown in Fig. S7C, if $\tau_{\mathrm{m}} \gg \tau_{\mathrm{v}}$, Eq. 23 predicts $\theta_\infty = \lim_{t\to\infty}\theta = \pi/2$, corresponding to complete coalescence. In contrast, if $\tau_{\mathrm{m}}$ is smaller than or comparable to $\tau_{\mathrm{v}}$, Eq. 23 predicts $\theta_\infty < \pi/2$, corresponding to incomplete coalescence.

**Cyst fusion modeling.** Here we rationalize the experimentally fitted nucleation rates pertinent to hole opening during cyst fusion. The nucleation rate is:

$$\Gamma \sim \mathrm{e}^{-\frac{\beta\pi\mathcal{T}^2}{\sigma}}, \tag{24}$$

where $\beta^{-1}$ quantifies the magnitude of the active noise, $\mathcal{T}$ is the line tension opposing hole formation, and $\sigma$ the monolayer tension.

Hole opening is driven by tension in a stretched cell monolayer, where cell stretching is comparable to that observed during micropipette aspiration, $L \approx 1.5L_{\mathrm{base}}$. We thus estimate $\sigma \approx \sigma_{\mathrm{asp}}$. Hole opening is resisted by the line tension $\mathcal{T}$, which for WT has been estimated in ablation experiments by Mukenhirn et al., $\mathcal{T}^{\mathrm{WT}} \approx 1$ nN (6). To obtain a rough estimate of KO line tension value, we consider that the evolution of the hole radius $r$ is described by the scaling equation (11)

$$\sigma - \frac{\mathcal{T}}{r} \sim \eta_{\mathrm{s}} r \frac{\mathrm{d}r}{\mathrm{d}t}, \tag{25}$$

where the left-hand side represents the net driving force and the right-hand side represents the viscous dissipation. Here we expect $\sigma \approx \sigma_{\mathrm{asp}}$, since cell stretching is comparable to that in micropipette aspiration. Experimentally, we observe that the same short-time dynamics of hole opening for WT and KO cysts (Figs. 2.J and 4.F in the main text), which suggests that both ratios $\sigma_{\mathrm{asp}}/\eta_{\mathrm{s}}$ and $\mathcal{T}/\eta_{\mathrm{s}}$ are comparable between WT and KO. Equality of the first ratio is indeed demonstrated by our micropipette experiments. Equality of the second term suggests that $\mathcal{T}^{\mathrm{KO}} \approx \mathcal{T}^{\mathrm{WT}}\eta_{\mathrm{s}}^{\mathrm{KO}}/\eta_{\mathrm{s}}^{\mathrm{WT}}$, leading to the estimate $\mathcal{T}^{\mathrm{KO}} \approx 0.5$ nN.

The required nucleation size to initiate hole opening corresponds to a zero net driving force in Eq. 25, i.e., $r_{\min} = \mathcal{T}/\sigma_{\text{asp}} \approx 0.1$ µm. In contrast, opening a hole in an isolated spherical cyst would require the nucleation of a much larger initial hole, $r_{\min} = \mathcal{T}/\sigma_{\text{base}} \approx 2$ µm, consistent with the fact that spontaneous hole opening in isolated cysts is not observed experimentally.

To assess the consistency of fitted $\Gamma$ values in the main text, we plot the energy barrier, $\mathcal{T}^2/\sigma$, versus the time required to nucleate a hole, $\Gamma^{-1}$. As shown in Fig. S7.D, our data align well with the theoretical prediction. Despite uncertainties in estimating $\mathcal{T}$ and $\sigma_{\text{asp}}$, both our data and published measurements for MDCK monolayers under stretch (1) are consistent with this theoretical framework.

**Movie S1.** **MDCK WT cysts doublets coalescence starting at day 2 after seeding in cavities. ZO1-mNeonGreen in green, H2B-mCherry in magenta, Z-projection, scale bar 20 µm, time in hh.**

**Movie S2.** **MDCK WT cysts doublets lumen fusion starting at day 4 after seeding in cavities. ZO1-mNeonGreen in green, H2B-mCherry in magenta, central slice, scale bar 20 µm, time in d:hh:mm.**

**Movie S3.** **Micropipette aspiration of an MDCK WT cyst (day 4) to measure the critical difference in pressure. This measurement is used to determine the monolayer tension $\sigma_{\mathrm{asp}}$. Scale bar 20 µm, time in mm:ss.**

**Movie S4.** **Micropipette aspiration of an MDCK WT cyst (day 4) to measure the aspired length dynamics. This measurement is used to determine the viscosity $\eta$. Scale bar 20 µm, time in mm:ss.**

**Movie S5.** **Laser cutting of an MDCK WT cyst (day 4) to measure the initial baseline tension $\sigma_{\mathrm{base}}$ and long term surface tension $\sigma_{\mathrm{cut}}$. Scale bar 20 µm, time in mm:ss.**

**Movie S6.** **MDCK WT and E-cad KO (H2B-mNeonGreen) cysts interaction starting at day 2 after seeding in cavities. WT cyst is labelled with ZO1-mNeonGreen in green and H2B-mCherry in magenta and the E-cad KO cyst with H2B-mNeonGreen in green. Cells from the E-cad KO cyst move around the WT cyst which is consistent with the tension measured by micropipette aspiration ($\sigma_{\mathrm{asp,WT}} > \sigma_{\mathrm{asp,KO}}$). Z-projection, scale bar 20 µm, time in hh.**
